# Cross-region neuron co-firing mediated by ripple oscillations supports distributed working memory representations

**DOI:** 10.1101/2025.09.04.674061

**Authors:** Ilya A. Verzhbinsky, Jonathan Daume, Sophia Cheng, Ueli Rutishauser, Eric Halgren

## Abstract

High-frequency (∼90Hz) ripple oscillations may promote integrative processing in mammalian brains. Previous work has demonstrated that the co-occurrence of these ripple oscillations is associated with enhanced temporal binding of neural activity between nearby human cortical neurons. However, it remains unclear whether such co-ripple facilitation of neuronal coupling supports cognitive processing, or if it occurs at greater distances. Here, we analyze intracranial recordings from patients implanted with microwire electrodes in the hippocampus, amygdala, ventromedial prefrontal cortex, anterior cingulate cortex, and pre-supplementary motor area, bilaterally, during a working memory task. We demonstrate that ripple oscillations significantly increase in all recorded regions, during encoding, maintenance and retrieval. Furthermore, co-occurrence of ripples increases between brain regions, associated with ∼30% increases in cross-region co-firing, both without decrement over distances up to 220 mm. These increases in cross-regional co-rippling and associated co-firing scale with memory load during maintenance and retrieval. During retrieval, co-ripples promote the reinstatement of the stimulus-specific long-distance co-firing patterns observed during encoding, especially during rapid recognition. Our findings reveal that co-occurring ripple oscillations orchestrate long-range neural communication that supports distributed neural representations during human cognition.

## INTRODUCTION

The ability of distant brain regions to communicate effectively is fundamental to cognition, but the mechanisms facilitating long-range neuronal integration during cognitive processing remain incompletely understood. High frequency synchronous oscillations have been proposed to assist in such integration during complex cognitive operations [1–5] such as working memory (WM), but their characteristics in humans have not been fully delineated [6], and their role remains the subject of considerable debate [7].

High-frequency ripple oscillations, first identified as components of hippocampal sharp wave-ripple complexes in rodents, have traditionally been associated with memory consolidation during sleep and quiet wakefulness [8, 9]. While initially discovered embedded within sharp waves in the hippocampus, subsequent research has demonstrated that similar oscillations occur widely throughout the cortex, often independently of any sharp wave activity [10–15]. These brief (50-100 ms) bursts of oscillatory activity facilitate the replay of previously experienced sequences of neuronal firing in rodents [16, 17], suggesting a role in information transfer between hippocampus and neocortex [18, 19]. Furthermore, the widespread detection of these ripple oscillations across various cortical regions in both awake animals [20, 21] and humans [11, 13–15, 22–26] indicates that they represent a general cortical phenomenon rather than a hippocampus-specific event, raising the possibility that ripples might coordinate neuronal activity across widely distributed brain networks during active cognition by transiently synchronizing neural activity across distant neural populations.

Recent investigations in humans have demonstrated that neural firing patterns are coordinated with local cortical ripples during spontaneous behavior [26], memory retrieval [22] and visual categorization [15]. However, critical questions remain unresolved. First, it is unknown whether ripples coordinate neural firing across brain-wide spatial scales, rather than only locally. Second, and more fundamentally, it has not been established whether ripple-mediated co-firing is systematically related to cognitive demands — scaling with task difficulty, carrying stimulus-specific information, and predicting behavioral performance. Addressing these questions requires simultaneous single-unit and LFP recordings across multiple brain regions during a cognitive task, a combination that has not previously been available. To fill this gap, we analyzed an open-access dataset of simultaneous single-unit and local field potential (LFP) recordings from intracranial Behnke-Fried microwires implanted in patients undergoing diagnostic monitoring for intractable epilepsy. We examined if co-occurring ripples across limbic and frontal areas facilitate the temporal coupling of neuronal firing during a Sternberg working memory task [27]. This large dataset, which contains widespread unit spike trains and LFP recordings, makes it possible to directly examine for the first time whether synchronous ripples in different forebrain locations and hemispheres provide a physiological environment conducive to greater interaction of neuronal firing between those locations, and whether those co-ripples and coordinated unit-firing are systematically related to task demands.

We report three main findings that go beyond what could be established from prior LFP-only or short-range unit recordings. First, co-occurring ripples between distant brain regions (up to 220 mm) are associated with ∼30% increases in cross-region co-firing that reflect genuine temporal coordination beyond independent rate increases. Second, both co-rippling and associated co-firing scale with memory load during maintenance and retrieval, establishing a link between ripple-mediated neural coupling and cognitive demands not previously shown at the single-neuron level. Third, and most critically, co-ripples during retrieval promote the reinstatement of stimulus-specific co-firing patterns observed during encoding, particularly during rapid recognition — demonstrating for the first time that co-ripples carry content-specific neural representations that predict behavioral efficiency, rather than merely reflecting elevated excitability. These findings support ripple-mediated neural synchronization as a general mechanism for cognitive integration across the human forebrain. Critically, the inclusion of single-unit recordings reveals the stimulus-specific informational content of this coordination — a dimension that LFP measures alone cannot access. By bridging oscillatory dynamics with single-neuron representations, our data establish co-ripple-mediated co-firing as a physiological substrate for distributed cognitive processing in the human brain.

## RESULTS

### Ripples and units are detected in all areas during a Sternberg working memory task

We analyzed intracranial recordings from a dataset previously published by Daume et al. [27]. In our study, we included data from 35 patients across 43 sessions (14 males, 21 females; age range 20-67 years) implanted with electrodes for evaluation of medically refractory epilepsy (Fig. 1A,B). Electrodes targeted the ventromedial prefrontal cortex (vmPFC), anterior cingulate cortex (ACC), pre-supplementary motor area (preSMA), amygdala (AMY), and hippocampus (HIP), bilaterally. Local field potentials (LFPs) and single-unit activity were simultaneously recorded and subsequently analyzed from each of the 8 microwires. Single units were identified based on previously reported methods [26, 28], and single units were merged if they were simultaneously detected on two microwires within the same bundle (see Methods). Patients performed a modified Sternberg working memory task (Fig. 1B) with varying memory loads (1 or 3 items), during which they were instructed to remember a set of images presented sequentially (encoding phase), maintain the memory over a delay period (maintenance phase), and identify whether a probe image was part of the original set (retrieval phase). A total of 1373 single units were isolated and classified across 1927 microwire channels (Fig. 1C). Ripples were detected in each brain region (Fig. 1D,E) using established methods [10] (peak of 70-100 Hz band-filtered LFP exceeding 2.5 standard deviations above mean power, at least 3 oscillations, the absence of sharp transients; see Methods. Care was taken to exclude epileptiform periods or electrodes implanted in the seizure onset zone). Per-region mean ± S.E.M ripple density, amplitude, duration, and frequency (Fig. 1F-I) were as follows: hippocampus (0.53 ± 0.09 sec⁻¹, 9.89 ± 9.00 μV, 73 ± 14 ms, 90.7 ± 1.3 Hz), amygdala (0.52 ± 0.09 sec⁻¹, 9.24 ± 8.51 μV, 75 ± 13 ms, 90.9 ± 1.1 Hz), vmPFC (0.53 ± 0.09 sec⁻¹, 7.23 ± 5.14 μV, 72 ± 11 ms, 91.1 ± 0.9 Hz), ACC (0.51 ± 0.09 sec⁻¹, 5.85 ± 3.39 μV, 69 ± 8 ms, 91.3 ± 0.7 Hz), and preSMA (0.55 ± 0.09 sec⁻¹, 6.92 ± 3.93 μV, 72 ± 10 ms, 91.2 ± 0.9 Hz). Ripple characteristics showed consistent profiles across patients despite differences in precise electrode locations (Supplementary Fig. 1). When a ripple was detected on any microwire channel within a bundle, single units recorded from that bundle fired an average of 0.23 ± 0.26 action potentials during the ripple event. No comparisons were made between units recorded on the same microwire to avoid contamination due to imperfect sorting.

**Figure 1.**
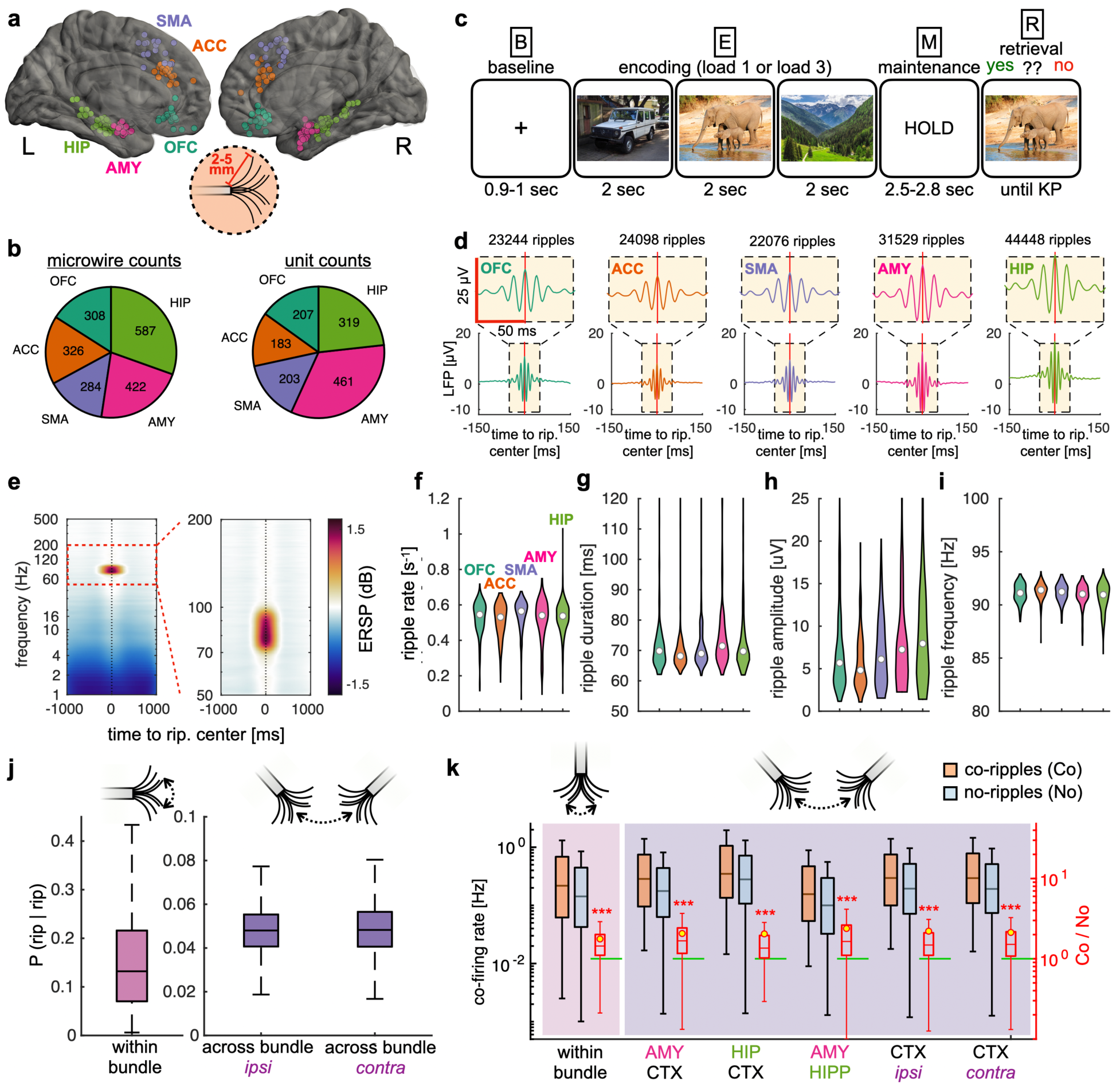
Microwire locations, task, ripple characteristics, and increased co-firing during co-ripples at all recording separations. **(a)** Electrode locations in MNI152 coordinates overlaid on a template brain. Both left (L) and right (R) hemispheres are shown. Inset shows a schematic of typical microwire electrode spray. Abbreviations: OFC, orbitofrontal cortex (ventromedial prefrontal cortex recording sites); ACC, anterior cingulate cortex; SMA, pre-supplementary motor area; AMY, amygdala; HIP, hippocampus **(b)** Number of electrodes and single units recorded from each region. **(c)** Overview of the Sternberg task. After a baseline period (B), either one or three (load 1 / load 3) images (E1,E2,E3) are shown separated out by a variable blank screen for 0.17-200 ms, followed by a 2.5s maintenance period (M). The participant is then shown a retrieval image (R) and responds with a keypress (KP) to indicate if they recognize the photo from the previous set. Images shown here are from the same categories as the set used in the task, but are not identical due to copyright. **(d)** Average broadband (0.1 – 1000 Hz) LFP locked to ripple centers in all 5 regions. **(e)** Average time-frequency plot across all microelectrodes locked to ripple centers (left), and zoomed into the high gamma range (right). **(f-i)** Distribution of per-channel ripple characteristics separated by recording location. Data is plotted for **(f)** event density, **(g)** duration, **(h)** amplitude and **(i)** oscillation frequency. Circle shows median and violin plots show the underlying distribution. **(j)** Conditional probabilities of ripple co-occurrences (< 25 ms overlap) between channels. Data is shown across microwires within the same bundle (left) and between different bundles within and across hemispheres (right). **(k)** Co-firing rate during co-ripples and no-ripples. Filled-in box plots denote median and inter quartile intervals, while whiskers denote 95% CI. Red outlined box plots to the right of each Co and No pair show distribution of the ratio between co-ripple (Co) and no-ripple (No) co-firing. Yellow circles are at the mean and horizontal green line is at Co/No = 1. Data is shown for unit pairs within bundle (left) and across bundle (right). Across bundle data is separated across amygdala-cortex (AMY-CTX), hippocampus-cortex (HIP-CTX), amygdala-hippocampus (AMY-HIP), ipsilateral cortico-cortical (CTX ipsi) and contralateral cortico-cortical (CTX contra) bundle pairs. Statistics were performed using a median permutation test on the Co/No distribution. *** p < 0.0001

### Ripples co-occur across long separations without significant decrease with distance

To characterize the spatial properties of ripple-mediated neural coordination, we analyzed how ripple co-occurrence (i.e., the percent of ripples on two contacts that overlap by 25ms or more) and associated unit co-firing varied with distance between recording sites. We first examined ripple co-occurrence within microwire bundles (intra-bundle, <∼4 mm separation) and found rates comparable to previous reports at the microscale [26], with a median co-occurrence rate of 13% [IQR 7%, 22%] within a bundle.

When examining ripple co-occurrence across greater distances, we found that while co-occurrence rates were lower between bundles (5% [IQR 4% 6%]) compared to within bundles, there was minimal further reduction as a function of fiber tract length between microwire bundles. Between different microwire bundles within the same hemisphere (inter-bundle, 71-203 mm separation), median ripple co-occurrence rates were 5% [IQR 4% 6%] (Fig. 1J). Surprisingly, cross-hemispheric (35-223 mm separation) ripple co-occurrence rates were almost identical (Δ 0.1% co-occurrence rate between intra and cross hemisphere co-ripple rates), indicating that once ripples extend beyond the local bundle, their co-occurrence probability remains relatively constant regardless of distance.

### Single neuron firing co-occurs preferentially during co-ripples

To determine whether co-ripples facilitate neural communication between distant brain regions, we analyzed unit co-firing (spikes co-occurring within a 25 ms window) during periods with co-occurring ripples versus periods without ripples. Specifically, we examined whether co-ripple periods were associated with enhanced co-firing relative to no-ripple periods, and whether this enhancement varied across different anatomical connections and distances.

We compared co-firing rates between units detected on wires within the same bundle and co-firing between units detected on different bundles. For each cell-pair, co-ripple periods were defined as times when any microwire in both units’ respective bundles contained detected ripples. Conversely, no-ripple periods were defined as times when no ripples were detected in either bundle.

Across all analyzed unit pairs (n = 31489), we observed significantly higher co-firing rates during co-ripple periods compared to no-ripple periods (median increase 34% [IQR-2 – 163%], p = 0, paired permutation test). We also found that co-firing during co-ripples was comparable between units recorded from the same microwire bundle (median co-fire rate 0.18Hz [IQR 0.04–0.62Hz]) and units recorded from different bundles (median co-fire rate 0.16 Hz [IQR 0.03–0.53Hz]), p = 0.001, permutation test of difference between medians). While the difference between local and distant co-firing was significant, the effect size was minimal, challenging the intuitive expectation that co-firing would be stronger for nearby neurons.

To further investigate how ripple-mediated coordination varies across specific anatomical pathways, we categorized across-bundle unit pairs into five connection types: amygdala-cortex (AMY-CTX, n = 1382 pairs), hippocampus-cortex (HIP-CTX, n = 210 pairs), amygdala-hippocampus (AMY-HIP, n = 3119 pairs), ipsilateral cortico-cortical (CTX ipsi, n = 2216 pairs), and contralateral cortico-cortical (CTX contra, n = 3748 pairs). All five connection types showed significant enhancement of co-firing during co-ripple periods (coR) compared to no-ripple periods (noR) (Fig. 1K). The magnitude of this enhancement was broadly similar across connection types, with median increases of 49% [IQR: 9 – 160%, p = 0, paired permutation test, coR vs noR] for AMY-CTX, 21% [IQR: 2 – 113%, p = 0] for HIP-CTX, 43% [IQR: 6 – 166%, p = 0] for AMY-HIP, 42% [IQR: 8 – 120%, p = 0] for CTX ipsi, and 44% [IQR: 5 – 140%, p = 0] for CTX contra pairs.

Next, we examined how the magnitude of co-firing enhancement between co-ripple and no-ripple periods varied with fiber tract distances between recording sites. Surprisingly, we found that the difference between co-ripple and no-ripple co-firing increased slightly with distance (r = 0.04, p = 5.3e-14). To ensure these findings were not driven by differences in baseline firing rates or recording quality across regions, we normalized co-firing rates by the mean of the individual firing rates of each unit pair, which yielded similar trends as a function of distance (Supplementary Fig. 2).

To determine whether elevated co-firing during co-ripples reflects true interaction greater than would be expected as a statistical consequence of independent increases in firing rate, we performed two rate-corrected analyses. First, we computed the observed co-firing rate within a ±25 ms coincidence window relative to the rate expected from independent firing (expected = rate_A_ × rate_B_ × window × 2), calculating individual firing rates separately for co-ripple and no-ripple periods. Co-firing exceeded the expected null during both conditions (co-ripples: 0.059 ± 0.001 Hz above null, p ∼ 0; no-ripples: 0.038 ± 0.001 Hz above null, p ∼ 0), but co-ripples showed a 56% greater elevation than no-ripples (p = 1.1e-99, paired t-test). We further validated this finding using the Spike Time Tiling Coefficient (STTC), a metric that quantifies spike timing coordination while explicitly correcting for firing rate[29]. STTC was significantly above zero during both co-ripple (0.023 ± 0.0009, p = 1.4e-136) and no-ripple periods (0.011 ± 0.0003, p = 1.4e-290), but was 117% greater during co-ripples (p = 1.7e-53, paired t-test, n = 26005 unit pairs), confirming that co-ripples enhance temporal coordination beyond what would be expected from elevated excitability alone (Supplementary Fig. 4). These findings reveal that ripple oscillations and neuronal firing are coordinated across widely distributed brain networks with minimal attenuation over anatomical distance. This distance-invariant enhancement of communication provides a potential mechanism for binding distributed neural representations during cognitive processing.

### Ripple and unit-firing rates increase in all regions during all task phases

To characterize the dynamics of neural activity during different phases of the working memory task, we quantified ripple occurrence rates and associated unit firing across all recorded brain regions. We observed systematic modulation of ripple activity that varied across task phases, with distinct patterns in limbic (hippocampus, amygdala), and frontal (preSMA, vmPFC, ACC) regions. During the encoding and maintenance phases, ripple rates increased moderately compared to pre-trial baseline levels in all recorded regions (encoding: 13% increase, maintenance: 10% increase, Fig. 2A). The greatest increase in ripple rates relative to baseline during encoding was observed in the hippocampus (20% increase, p < 1e-10, permutation test) and amygdala (16% increase, p < 1e-10), while the greatest increase during maintenance was observed in the vmPFC (13% increase, p < 1e-10) and hippocampus (13% increase, p < 1e-10). The task-related modulation of ripple activity was most pronounced during the retrieval phase, where we observed substantial increases in ripple rates across hippocampus (13% increase from baseline, p < 1e-10), amygdala (12% increase, p < 1e-10), and preSMA (36% increase, p < 1e-10).

**Figure 2.**
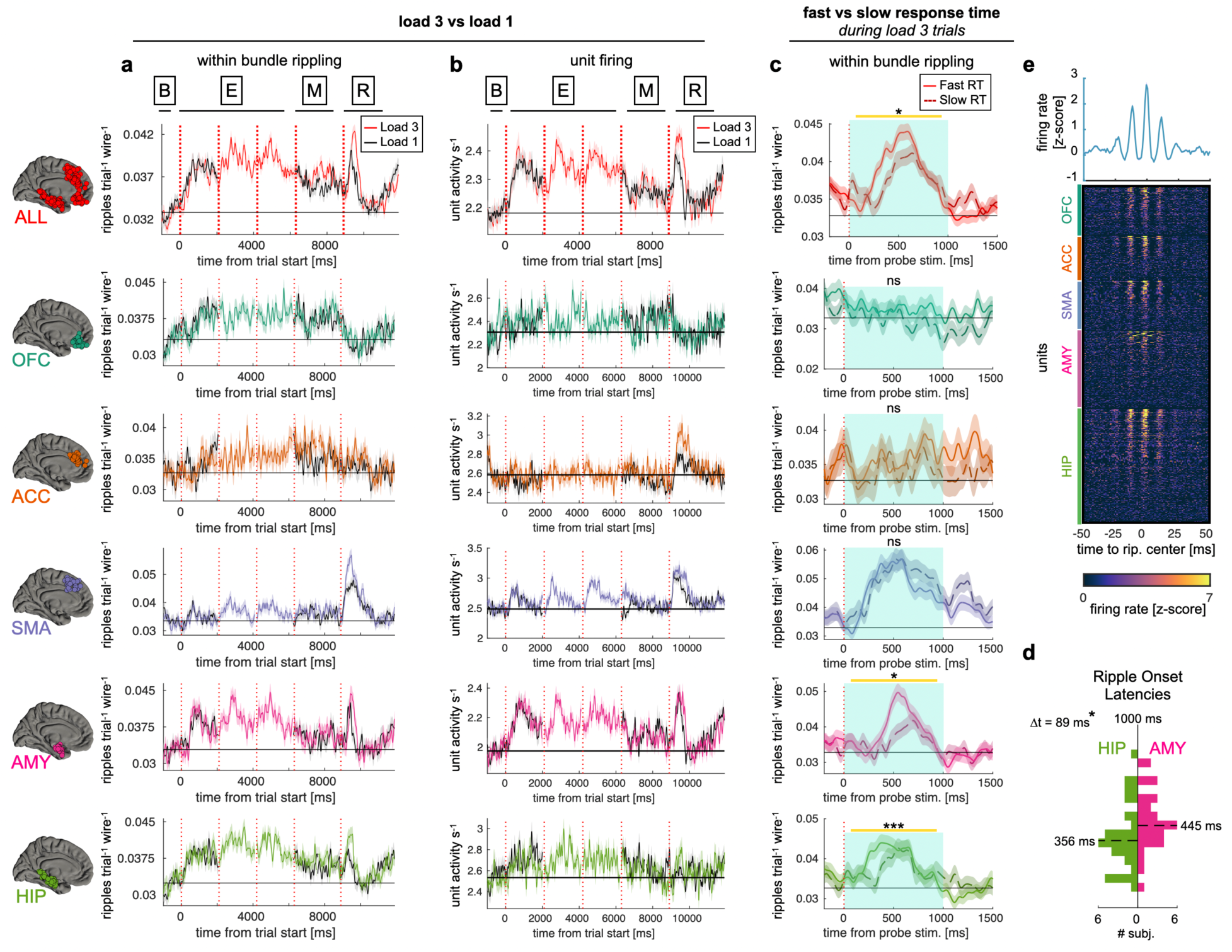
Task modulation of ripples and units within regions according to task variables. **(a)** Ripple oscillation rates within each sampled region. Data is shown for mean number of ripples per electrode within a given bundle across all trials. Vertical dashed lines show the beginning of each task stage (encoding, maintenance, retrieval). Colored curve shows mean ± sem for all load 3 trials, while black curve shows load 1 trials. A gap is shown in the maintenance period because its duration is variable. For plotting purposes, data are smoothed with a 100 ms sliding gaussian window. **(b)** Same as **(a)** but for mean firing rate across all detected single neurons within each region. **(c)** Ripple oscillation rates for fast (solid line) and slow (dashed line) responses after the retrieval stimulus is shown in load 3 trials. To control for the uneven distribution of microwire yield across each patient, data is shown as mean number of ripples within each bundle *per microwire* per trial. Statistics were performed during the turquoise shaded 0- to 1000-ms interval following retrieval stimulus presentations. Statistics for load 1 vs load 3 in **(a)** and **(b)** are shown in Supplementary Figure 3. * p < 0.01 *** p < 0.0001, ns, not statistically significant. linear mixed-effects models with patient as random effect. **(d)** Ripple onset latency distribution following retrieval probe. Distribution is shown across subjects for amygdala and hippocampus electrodes. **(e) (Top)** Population-averaged firing rate aligned to ripple center (time = 0 ms), showing rhythmic modulation of spiking activity consistent with phase-locking to ripple oscillations. **(Bottom)** Firing rate heatmap for all recorded units aligned to ripple center. Units are sorted by region followed by peak firing rate. Color scale indicates instantaneous firing rate (z-scored for each cell on the raster centered ± 3 seconds around ripples centers). The majority of units show transient increases in firing rate time-locked to ripple events, with visible oscillatory structure reflecting spike-phase coupling to the underlying ripple rhythm.

Importantly, mean activity across all detected single units activity demonstrated similar, although not identical, modulation patterns (Fig 2B, see Supplementary Fig. 5 for analysis across subjects), with neuronal firing rates showing significant but modest positive correlations with ripple rates during encoding (r = 0.21, p = 1.8e-115), maintenance (r = 0.21, p = 1.41e-59) and retrieval phases (r = 0.25, p = 1.9e-87) in all regions. Overall firing increased by 6% during encoding, 4% during maintenance, and 6% during retrieval. Within-ripple firing rates of neurons also increased significantly during these phases compared to ripples occurring during baseline periods (2.95 Hz vs 2.83 Hz, p = 4.30e-3, paired permutation test), suggesting enhanced neuronal recruitment during task-relevant ripples. Consistent with prior work [11, 22, 23, 26], alignment of single unit firing to ripple centers revealed rhythmic modulation of spiking activity consistent with phase-locking to the underlying ripple oscillation across all regions (Fig. 2e), confirming that detected events reflect genuine oscillatory coordination of local neural populations.

### Co-ripple rates and engagement of units during co-ripples scale with WM load and WM processing efficiency

Within region co-ripple rates and unit firing rates were also differentially modulated by memory load. Co-ripple rates increased during maintenance according to memory demand across the entire set of microwires (p = 0.038, load 3 vs. load 1, FDR-corrected linear mixed effects) and within ACC (p = 0.006). Firing rates across all neurons did not increase significantly with memory demand during maintenance (p=0.99), but did increase within the subset of neurons located in the ACC (p = 0.03) and preSMA (p = 0.003). During the retrieval phase, co-ripple rates also increased with memory load across all microwires (p = 3.2e-6) as well as within preSMA (p = 0.007), AMY (p=0.04) and HIP (p=7.8e-5). Similarly, firing rates across all neurons increased with memory demand during retrieval (p=0.001), as well as for neurons located in the ACC (p = 1e-4), preSMA (p = 0.002) and AMY (p=0.03). This systematic scaling of co-ripple rates with memory load — observed during both maintenance and retrieval — directly links ripple-mediated coordination to cognitive demand, establishing that co-ripples do not simply reflect a fixed physiological process but are dynamically recruited in proportion to the computational requirements of the task.

We next investigated the relationship between ripple activity and behavioral performance, specifically whether there was any relationship between ripple rates during fast and slow response times (RTs). We only analyzed rippling in high memory load trials (i.e., load 3), defining fast and slow responses as above and below the median response time for load 3 trials. We found that overall ripple rates after the retrieval stimulus were significantly higher during fast response trials compared to slow response trials, specifically in the hippocampus (p = 9.3e-4, linear mixed effects) and amygdala (p = 0.013). This association between ripple rates and response speed suggests that the efficiency of working memory retrieval is predicted by the degree of ripple engagement — a relationship paralleling the fast/slow dissociations that have been central evidence for the involvement of specific neural mechanisms in working memory processing.

Temporal analysis revealed that hippocampal ripple onsets consistently preceded the amygdala onsets in both fast and slow response conditions (hippocampus: 379 ± 33 ms; amygdala: 473 ± 30 ms; two sample t-test, p = 0.041), indicating that hippocampal ripples temporally lead amygdala ripples during memory retrieval.

These findings indicate that ripple oscillations and unit firing in both limbic and frontal regions are dynamically modulated during working memory processes, with robust activation during all phases of the task, especially when memory load is higher. These findings indicate that not only the magnitude but also the timing of ripple activity, particularly in limbic structures, is associated with efficient working memory processing.

### Co-ripples are most related to stimulus but not motor processing

Beyond the stimulus-locked, load modulated and response timing effects that form the core of our findings, we examined multiple task-related parameters to comprehensively characterize ripple and unit dynamics during working memory processing. Specifically, we investigated ripple rates when retrieval stimuli matched versus mismatched the encoded stimulus set (Supplementary Fig. 6a). We also attempted to examine task accuracy (Supplementary Fig. 6b), but error trials were not analyzed due to the high accuracy of task performance across participants (mean accuracy: 93%), which resulted in insufficient incorrect trials for reliable statistical comparison. Finally, we analyzed neural activity locked to the response key press timing (Supplementary Fig. 7). While all of these analyses revealed significant modulation of both ripple oscillations and unit firing, we found that neural responses were most robustly engaged by stimulus-related processing compared to motor response execution. Furthermore, memory load manipulations elicited stronger and more consistent differences in ripple and unit activity profiles across brain regions than other task parameters. Given the superior signal-to-noise ratio observed in stimulus-locked analyses under varying memory loads and response times, we focused much of our investigation on these conditions, which provided the clearest window into understanding how ripple oscillations coordinate distributed neural communication during working memory processing.

### Long-range ripple co-occurrence selectively increases with memory load during maintenance and retrieval

Having established that ripple rates increase locally within individual brain regions during working memory, we next examined whether ripples co-occur across distant brain regions, potentially serving as a mechanism for long-range information integration. We defined co-occurring ripples (co-ripples) as periods where at least one ripple was simultaneously detected across two different microwire bundles. Across most possible region pairs, we observed a significant increase in ripple co-occurrence during both maintenance and retrieval stages of the task (Fig. 3). This effect was most pronounced between HIP-preSMA (48% change from baseline) during retrieval, and vmPFC-HIP (21% change from baseline) during maintenance.

**Figure 3.**
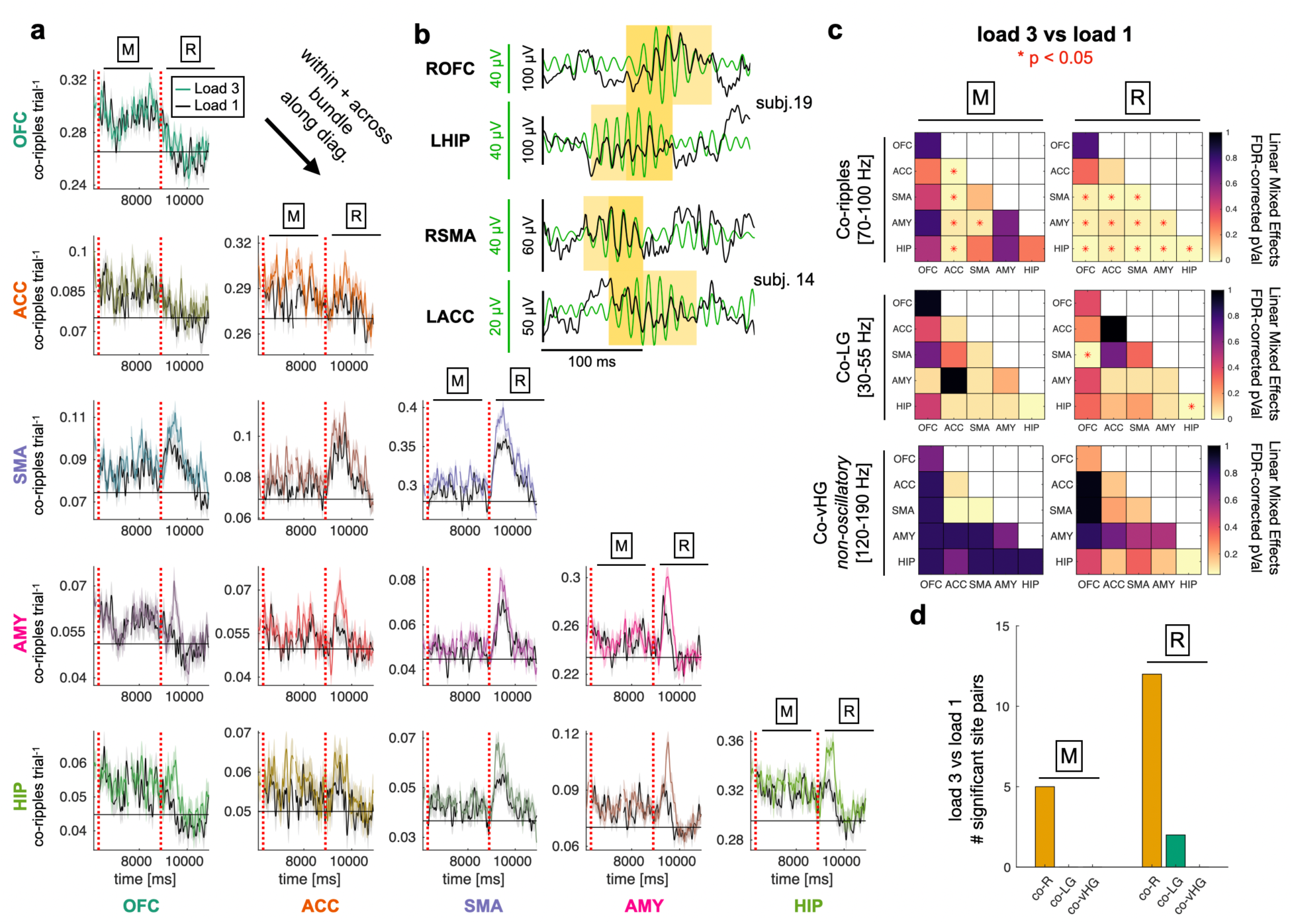
Widespread ripple co-occurrence increases with greater memory load. **(a)** Co-ripple rates between all recorded regions during maintenance (M) and retrieval (R) periods. Data is plotted as mean±SEM for each pair of locations with load 3 trials in color and load 1 in black. Diagonal shows across combined with within bundle data. Black horizontal line shows mean baseline co-ripple rate for each region pair. **(b)** Example of simultaneous ripple events detected across four brain regions in two patients. Raw LFP traces (black) and ripple-band filtered signals (70–100 Hz; green) are shown for right orbitofrontal cortex (ROFC), left hippocampus (LHIP), right supplementary motor area (RSMA), and left anterior cingulate cortex (LACC). Yellow shading indicates detected ripple events; darker shading denotes periods of co-ripple overlap across regions. Scale bars indicate voltage amplitude (green = filtered signal; black = raw LFP) and time (100 ms). **(c)** Results of linear mixed effects analysis of pairwise load 3 vs load 1 co-rippling across regions during maintenance and retrieval. Red stars indicate FDR-corrected p-values < 0.05 for detected co-ripples, low gamma (LG) co-oscillations and non-oscillatory very high gamma (vHG) co-events. **(d)** Summary of statistical analyses. Number of site pairs that show significant modulation during load 3 trials. Data is shown for co-ripples, co-LG and co-vHG. Note heightened response during maintenance and retrieval in the ripple band.

Critically, we also found that memory load significantly modulated the rate of co-ripples across brain regions. When participants maintained three items in working memory, the rate of co-ripples was substantially higher compared to trials with only one item (Fig. 3B,C). This load-dependent modulation was observed during both maintenance and retrieval phases, with stronger effects during retrieval (mean increase across all pairs: 2% increase from load 1 to load 3 during maintenance, p= 0.036, 6% increase during retrieval, p = 3.2e-6). The greatest significant increase in load 3 vs load 1 co-rippling during retrieval was observed between HIP-preSMA (18% increase, p = 0.001, FDR-corrected linear mixed effects) and vmPFC-HIP (17% increase, p = 3e-4). In contrast, load-modulated maintenance co-rippling was greatest between ACC-preSMA (8% increase, p = 0.01) and ACC-HIP (8% increase, p=0.03). Therefore, long distance co-rippling increases and is modulated by task load during maintenance and retrieval stages.

### Ripple co-occurrence shows greater modulation by memory load than other high gamma signals

To determine whether this load-dependent coordination was specific to the ripple frequency band, we performed parallel analyses on oscillatory events detected in low gamma (30-55 Hz) and very high gamma (120-190 Hz) frequency bands (see Methods). While we observed task-related increases in co-occurrence of oscillations in these other frequency bands, the contrast-to-noise (CNR) for load modulation was smaller during retrieval (low gamma: CNR = 6.21 between 1 to 3 WM items, p = 3.4e-4; very high gamma: CNR = 10.13, p = 1.5e-5) and maintenance (low gamma: CNR = 4.71, p = 8.0e-4; very high gamma: CNR = 2.70, p = 0.14) compared to the ripple band (retrieval: CNR = 12.42, p = 1.4e-7; maintenance CNR = 4.81, p = 8.0e-3, Fig. 3B,C, Fig S4C,D).

During the retrieval stage, linear mixed effects modelling revealed significant load-dependent modulation across 12 of the 15 site pairs during co-ripples, only 1 site pair during low gamma oscillations and 7 site pairs during very high gamma oscillations. Load-dependent oscillation co-occurrence during maintenance was also greatest for ripples (5 significant pairs) compared to low gamma (0 pairs) and very high gamma (2 pairs). These findings suggest that co-occurring ripples across limbic and frontal regions represent a specific neural mechanism for coordinating distributed brain activity during working memory, with the degree of coordination scaling with cognitive demand more than other high gamma signals. (Fig. 3B,C, Supplementary Fig. 8).

Unlike event-based co-ripple analyses, cross-regional amplitude envelope correlations did not show robust load-dependent modulation in any frequency band (Supplementary Fig. 9). While the ripple band showed numerically stronger load modulation than low gamma and very high gamma bands, only one region pair reached statistical significance. This suggests that the load-dependent coordination we observe is specific to the temporal co-occurrence of discrete ripple events rather than a general property of correlated high-frequency power.

### Ripple co-occurrence increases unit co-firing in each task stage

To investigate whether co-ripples facilitate neural communication between brain regions, we analyzed cross-regional unit co-firing during different task periods. We defined co-firing as the co-occurrence of spikes from units in separate brain regions within a 25 ms window. For each region pair, we calculated the percent change in co-firing rate relative to the median baseline co-firing rate during three conditions: 1) periods with co-occurring ripples (co-ripple periods), 2) periods without ripples in either region (no-ripple periods), and 3) across all periods regardless of ripple occurrence (all periods). In this analysis we only included unit pairs that exhibited a baseline co-firing rate of at least 0.025 Hz. This resulted in a 12335 out of a total 16274 cell pairs being included in the analysis.

Examination of unit activity revealed a robust increase in cross-regional unit co-firing specifically during co-ripple periods across all task stages (Fig. 4). During encoding, maintenance, and retrieval phases, co-ripple periods showed significant enhancement of median unit co-firing compared to baseline (encoding: 28% increase, p = 0, paired-permutation test of medians; maintenance: 29% increase, p = 0; retrieval: 24% increase, p = 0). This effect was most pronounced among cross-region pairs during maintenance between ACC-AMY (84% increase during co-ripples, p = 0) and preSMA-ACC pairs (86% increase, p = 0) during retrieval, but was consistently observed across almost all region pairs (Fig. 4A,C).

**Figure 4.**
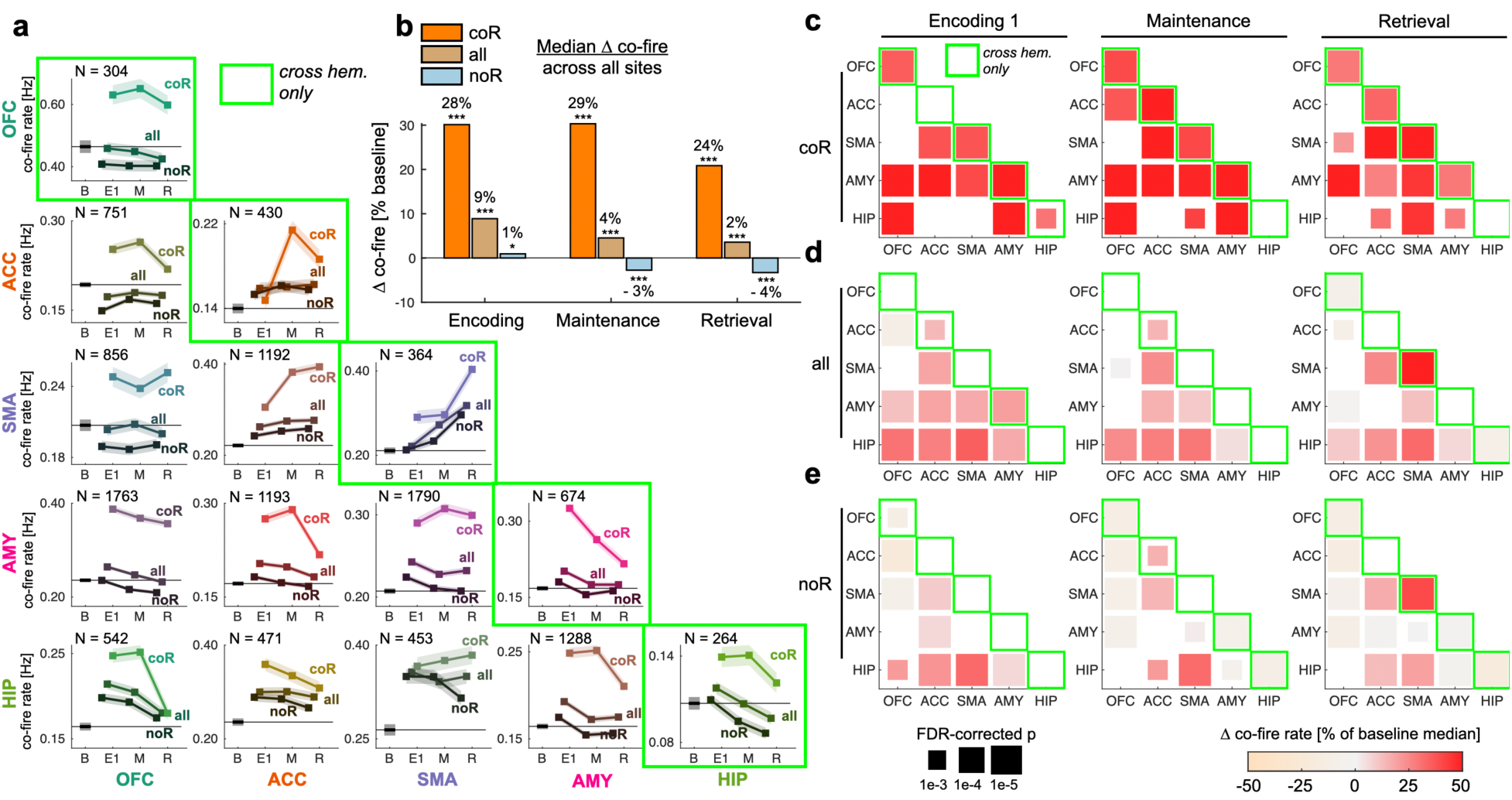
Task related modulation of cross-region neuron co-firing is mediated by ripple oscillations. **(a)** Inter-area co-firing rates during baseline (B), first image presentation (E1), maintenance period (M), and response retrieval (R). Co-firing is shown for all conditions (all), co-ripple periods (coR) and periods where neither site has a detected ripple (noR). Plots show median ± mean absolute deviation. Number of cell pairs is shown in the top left of each plot. Only cell pairs that have a co-fire rate >= 0.025 Hz are included in this analysis. **(b)** Distribution of inter-area co-firing across all recorded sites. Statistics are shown for each condition compared with baseline. Red stars indicate p < 0.05 FDR-corrected permutation-based test of difference between medians. **(c-e)** Co-firing rates vs baseline statistics during all task stages for coR **(c)**, all **(d)**, and noR **(e)** conditions. The color of each box indicates the magnitude of the difference from baseline, and the size of the box indicates the FDR-corrected significance (permutations test). The diagonal for **(a, c-e)** shows cross hemisphere co-firing between the same structure.

In contrast, no-ripple periods showed minimal to no change in co-firing rates compared to baseline across any task stage (encoding: 1% increase, p = 0.02; maintenance: 3% decrease, p = 0; retrieval: 4% decrease, p = 0, Fig. 4B). When examining all periods together regardless of ripple occurrence, we observed a modest increase in co-firing (encoding: 9% increase, p = 0; maintenance: 4% increase, p = 0; retrieval: 2% increase, p = 0, Fig. 4B), likely driven by the contribution of co-ripple periods. Overall, the difference between co-ripple, no-ripple and all period co-firing was significantly different during encoding (p = 1.73e-11, Kruskal-Wallis test), maintenance (p = 6.95e-82) and retrieval (p = 3.22e-80) stages. These results indicate that co-occurring ripples across brain regions specifically enhance the temporal coordination of neural firing, potentially providing a mechanism for information transfer between distant brain areas during cognitive processing.

### Unit co-firing is enhanced by memory load during co-ripples

Having observed that co-ripples enhance cross-regional unit co-firing throughout the task, we next examined whether this effect is modulated by memory load. For each region pair, we calculated the percent change in the amount of co-firing per trial between high memory load and low memory load conditions during co-ripple periods, no-ripple periods, and across all periods (see Methods).

Our analysis revealed a load-dependent enhancement of unit co-firing specifically during co-ripple periods (Fig. 5A). Across distributed region pairs, co-firing during co-ripples was substantially higher in the high load condition compared to the low load condition (maintenance: 13% increase from load 1 to load 3, p = 7.6e-39 paired Student’s t-test; retrieval: 19% increase, p = 4.38e-39). In contrast, no-ripple periods showed minimal modulation by memory load, with minimal to no significant increase in co-firing rates between high and low load conditions (maintenance: −1% change, p = 0.03; retrieval: −0.2% change, p = 0.77). When examining all periods together, we observed a modest load-dependent increase in co-firing (maintenance: 1% increase, p = 0.003; retrieval: 5% increase, p = 1e-15), substantially smaller than the effect observed during co-ripple periods.

**Figure 5.**
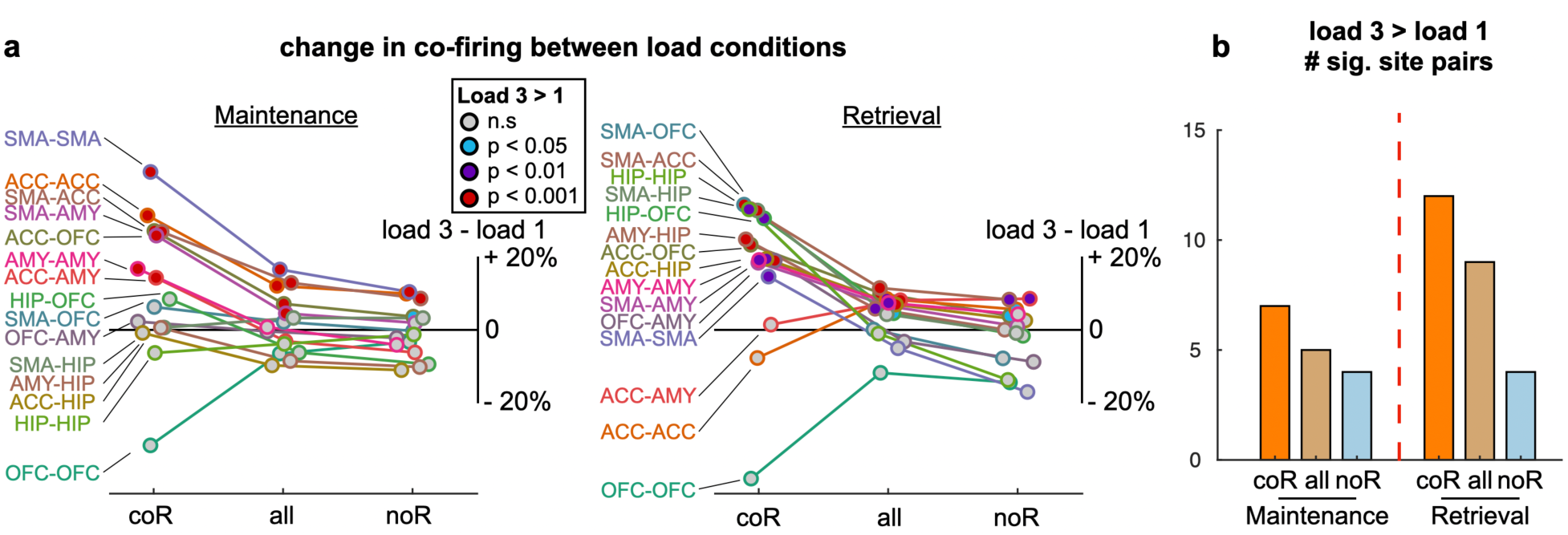
Co-ripple oscillations increase co-firing according to memory load. **(a)** Change in inter-area total co-firing for load 3 vs load 1 conditions in the maintenance, and retrieval periods, as a function of co-ripples between those areas. The percent change in co-firing is shown for all conditions (all), co-ripple periods (coR) and periods where neither site has a detected ripple (noR). Plots show mean percent change in co-firing for each site pair across all conditions. As in Fig 4, site pairs within the same structure (e.g., SMA-SMA) exclusively contain contralateral hemispheric microwires. Only cell pairs with a baseline co-firing rate of at least 0.025 Hz are analyzed. The color of the dots represents the significance of the load 3 > load 1 modulation, with p < 0.05 colored blue, p<1e-2 colored purple, and p < 1e-3 colored red, one-sided Student’s t-test, FDR-corrected. **(b)** Summary of statistics indicating the number of site pairs (out of 15 possible) whose co-firing is significantly elevated for load 3 compared to load 1 in Maintenance and Retrieval phases of the task.

The differential effect of memory load on co-firing during maintenance was significant in 7 site pairs during co-ripple periods, 3 site pairs during no-ripple periods and 6 site pairs during all periods of the recording. The effect was even greater during retrieval, with 13 significant site pairs during co-ripple periods, 3 site pairs during no-ripple periods and 11 site pairs during all periods (Fig. 5B). These findings suggest that co-ripples are specifically associated with the temporal window during which inter-regional communication is enhanced and dynamically modulated by cognitive demand, potentially serving as a mechanism for load-dependent scaling of information transfer during working memory processing.

### Co-ripples increase the repetition of stimulus-specific co-firing during retrieval

We next investigated whether co-ripples support the reinstatement of stimulus-specific neural representations across distant structures during memory retrieval. To this end, we examined the repetition of co-firing patterns between encoding and retrieval phases. For each cell pair recorded from different brain regions, we identified co-firing events (spikes within 25 ms) during the encoding phase when a specific stimulus was presented and during the retrieval phase when the same stimulus was presented (Fig 6A).

**Figure 6.**
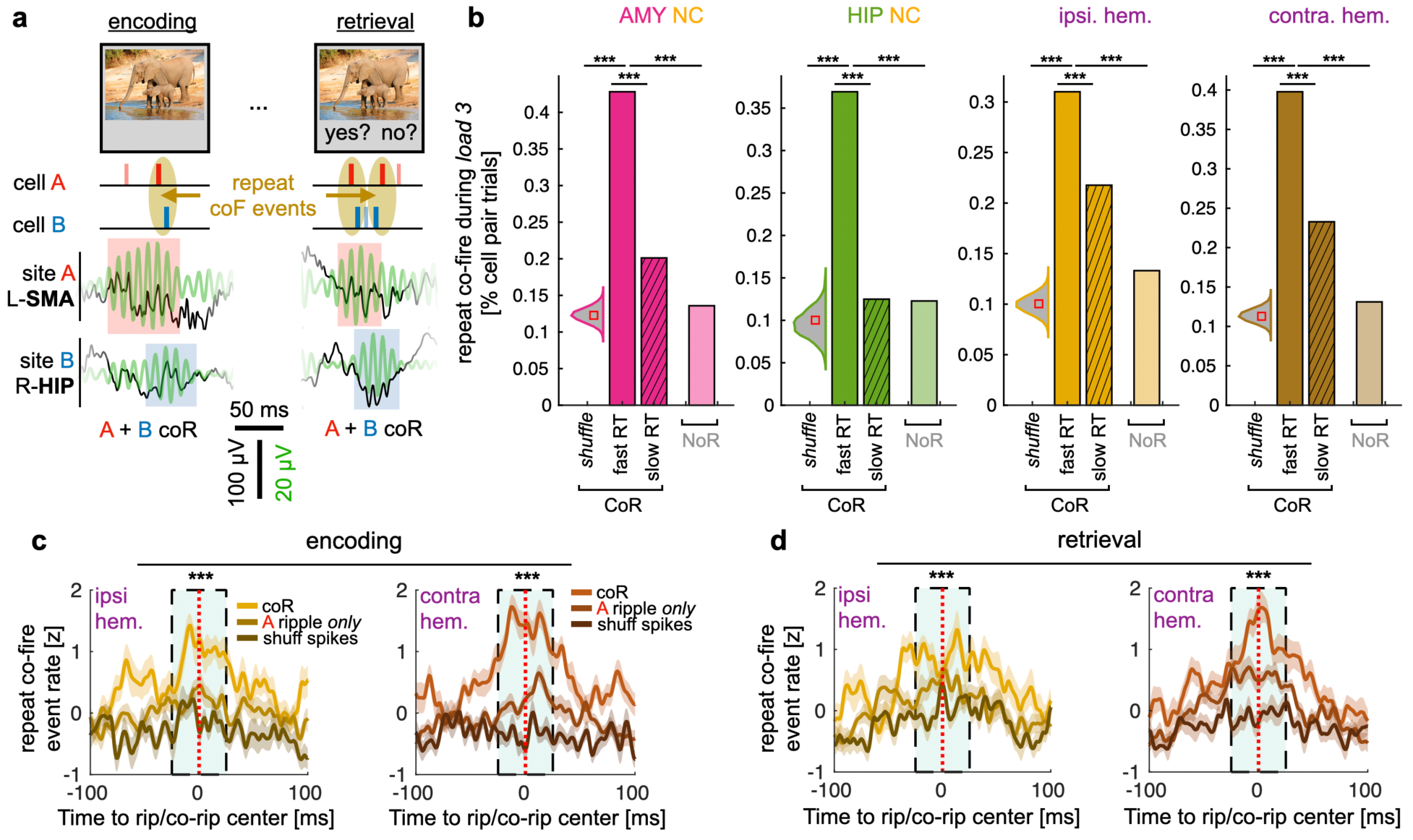
Increased repetition of stimulus-specific cross-structure firing during coR. **(a)** Example of repeated coF (co-firing) between encoding and retrieval phases on a trial when both sites co-rippled. For each cell pair across structures, we assessed whether they co-fired during encoding, and if so whether their co-firing was repeated during the retrieval when the same stimulus was presented. **(b)** Comparison of co-firing repetition during fast and slow response times in load 3 trials. Bars indicate the percentage of repeated co-firing across cell pairs and trials during the retrieval period when that co-firing pattern was previously observed for the same stimulus during the encoding period. Analysis shows co-firing repetition during co-ripples (**coR**) for trials with fast response times (below median RT) and slow response times (above median RT). Violin plots show the null distribution from a procedure where trial assignment of co-fire events are independently shuffled for encoding and retrieval stages (n = 1000 permutations, square shows median). Additionally, co-firing repetition rates are shown for all load 3 trials during duration-matched no-ripple periods (**noR**). Co-firing repetition during coR is significantly greater for fast responses compared to slow responses. All effects are replicated using amygdala-neocortical (AMY NC), hippocampus-neocortical (HIP NC) pairs, and when using all within hemisphere (ipsi hem.) or cross hemisphere (contra hem.) cell pairs (results for load 1 trials can be found in Supplementary Fig. 10c). **(c,d)** Time-course of repeated Encoding→Retrieval co-fire events relative to co-ripple centers. This is compared to when ripples occurred in site A only, and to shuffled spike times. Cyan box outlines the ± 25 ms window where the three conditions were compared. Repeated Encoding→Retrieval co-fire events increase during coR during both the **encoding (c)** and **retrieval stages (d)**, and again, these effects are replicated within hemisphere and cross hemisphere. * p < 0.05 *** p < 0.0001

We calculated the percentage of cell-pair trials (# cell-pairs × # total trials) that exhibited repeated co-firing patterns between encoding and retrieval phases. When we separated our analysis into co-ripple periods and no-ripple periods (duration-matched across 25 shuffles per cell-pair trial), we found that pattern repetition was substantially enhanced during co-ripple periods compared to no-ripple periods. During co-ripples, the repetition rate was 0.29%, while during no-ripple periods, this rate dropped to 0.14% (p = 0, χ² = 213.9). This enhancement during co-ripples exceeded what would be expected from the general increase in co-firing rates alone (Supplementary Fig. 10a).

The effect during co-ripples was present across different regional connections but was particularly pronounced for hippocampus-cortical pairs (0.28% during co-ripples vs 0.11% during no-ripples, p = 1.1e-16, χ² = 23.5) and amygdala-cortical pairs (0.31% during co-ripples vs 0.16% during no-ripples, p = 1.1e-16, χ² = 69.5). Effects were similar for cell pairs between ipsilateral (0.26% during co-ripples vs 0.14% during no-ripples, p = 0.002, χ² = 69.1) and contralateral regions (0.31% during co-ripples vs 0.14% during no-ripples, p = 0, χ² = 145.6). These effects were consistent at the individual subject level (Supplementary Fig. 10b).

Critically, we also found that co-ripple-mediated co-firing repetition was associated with behavioral performance (i.e., patient response time). We focused our analysis on high memory load (3 items) trials where the retrieval stimulus matched one of the encoded stimuli. This design ensured that the sensory input was identical during both encoding and retrieval phases, allowing us to isolate the contribution of behavioral variability while controlling for stimulus-related factors. The high-load condition also represented a more challenging task and provided the widest range of response latencies. During these high-load trials, co-firing repetition during co-ripples was significantly greater for trials with fast response times compared to slow response times (0.36% for fast responses vs 0.23% for slow responses, p = 3.3e-14, χ² = 57.5, Fig. 6B). Co-firing repetition effects during co-ripples were also present during low memory load (load 1) trials, though the relationship with reaction time was attenuated compared to high load conditions, consistent with reduced demands on cross-regional coordination during single-item retrieval (Supplementary Fig. 10c). These findings demonstrate that co-ripple-mediated repetition of neural patterns is associated with more efficient responses to the working memory retrieval.

We next determined the time-course of the reinstatement of stimulus-specific co-firing patterns during co-ripple activity, by plotting their occurrence relative to the centers of detected co-ripples (ripples occurring simultaneously in both recorded regions). We then tested the significance of co-firing enrichment around co-ripple centers compared to their enrichment around ripples that occur in only one of the two bundle locations. As an additional control, we created surrogate data by shuffling spike times within each trial while preserving the overall firing rate profiles and recomputed the enrichment of matching co-fire events around the original co-ripple centers. We found a marked increase in co-firing events around the centers of co-ripples compared to single ripple and shuffled spike controls (Fig. 6C,D). This increase was seen within hemispheres (p = 6.1e-8, one-way ANOVA) and across hemispheres (p = 7.2e-15) during the encoding stage, and within hemispheres (p = 4e-4) and across hemispheres (p = 3.2e-10) during the retrieval stage.

These findings demonstrate that co-ripples provide privileged temporal windows for repeating stimulus-specific neural patterns across distant brain regions during memory retrieval. Our results reveal an association between co-ripple-mediated neuronal firing repetition and behavioral performance, highlighting their functional importance in supporting efficient recognition memory. By facilitating the coordinated reactivation of encoding-related activity patterns, co-ripples may contribute to the neural processes necessary for successful working memory performance.

## Discussion

Our study provides the first direct evidence that co-occurring ripple oscillations coordinate stimulus-specific neuronal firing between distant human brain regions during a cognitive task. While previous work has established that co-ripples occur spontaneously across widespread cortical areas [10, 11], are modulated by task conditions at the LFP level [25], and facilitate local neuronal interactions at short cortical distances [15, 22, 26, 30], these studies could not determine whether co-ripples carry content-specific neural representations, whether their effects on neuronal co-firing are related to cognitive demand, or whether such coordination extends across brain-wide distances. By combining simultaneous single-unit and LFP recordings across limbic and frontal structures during a working memory task, we demonstrate that co-ripples enhance cross-region co-firing that is load-dependent, stimulus-specific, and predictive of behavioral efficiency — properties that could not be inferred from LFP measures or short-range unit recordings alone. Because neural computation ultimately depends on the spiking activity of individual neurons, directly examining how ripples shape the timing and content of cross-regional spike patterns is essential for moving beyond correlational LFP signatures toward a mechanistic understanding of how ripple synchrony mediates cognition.

Our most novel finding is that co-ripples during retrieval promote the reinstatement of stimulus-specific co-firing patterns observed during encoding, particularly during rapid recognition. Behavioral data from the Sternberg Paradigm was originally taken to support serial scanning of items being held in working memory, with subsequent work suggesting that in some circumstances a parallel interrogation of ‘memory strength’ is operative [31, 32]. Our finding that rapid reaction times are associated with repetition of a specific distributed firing-pattern evoked by a stimulus from its Encoding to its Retrieval may provide a mechanism whereby memory strength could be available on some trials. Specifically, multiple processes may be engaged by the Retrieval, and if reinstatement of firing patterns is successful, then a rapid response is made; otherwise slower (possibly sequential) processes support longer latency responses. Neurophysiological models of recognition judgments also often posit two parallel mechanisms: hippocampo-cortical firing-pattern reinstatement and facilitated processing from recent cortico-cortical activation [33]. Our observation of the former on faster trials suggests that one of the two processes active in working memory tasks is hippocampal-dependent, and that it may sometimes be responsible for quicker trials. Yonelinas [34] reviews the hippocampal contribution to working memory but proposes that it assists in familiarity rather than recollection.

A second key finding is that the degree of ripple-enhanced co-firing increases systematically with cognitive demand, with stronger coordination observed under high memory load conditions. This load-dependent modulation — demonstrated here at the single-neuron level for the first time — suggests that ripples provide a flexible mechanism for adjusting the strength of inter-regional communication according to task requirements. However, future work is needed to understand how ripples may coordinate with slower oscillations according to cognitive demand. An analysis of the same dataset included in this study by Daume and colleagues [27] demonstrated that theta-gamma phase-amplitude coupling (PAC) in human hippocampal neurons similarly scales with cognitive control demands, with PAC neurons showing enhanced phase-locking to frontal theta specifically during high-load conditions. This suggests that multiple oscillatory mechanisms — both ripples and cross-frequency coupling — work in concert (or perhaps in opposition [35]) to support flexible inter-regional communication during demanding cognitive tasks.

Importantly, the mechanisms supporting short-term working memory may be associated with broader memory related functions. In another study by Daume et al. [36], persistent activity of hippocampal neurons during working memory maintenance predicted successful long-term memory formation in a separate task, demonstrating a direct neuronal-to-neuronal link between working memory maintenance and long-term memory encoding across distributed networks. The convergent evidence from ripple-mediated coordination, phase-amplitude coupling mechanisms [27], and the shared neural substrates linking working and long-term memory systems [36] collectively support widespread-interactive models of cortical function, revealing multiple, complementary mechanisms through which the brain achieves flexible integration of information across its distributed networks.

The enhanced neural coordination we observed spans considerable physical distances, occurring between brain regions separated by white matter fiber tracts up to 220 mm, across different lobes, and even between hemispheres. Notably, neither co-firing nor the co-ripple-induced enhancement in co-firing decremented with distance, despite the well-established exponential decrease in cortico-cortical connectivity with distance [37, 38]. In prior studies without unit recordings, a similar lack of decrement with distance was found at long cortico-cortical separations for co-ripples in spontaneous sleep and waking [11], and at short separations for facilitation of firing co-prediction[26]. This distance-invariance suggests an emergent network phenomenon rather than simple point-to-point transmission of activity [1, 39]. While our sampling was necessarily limited to specific recording sites, the fact that robust co-ripples and associated co-firing were observed across diverse, unselected locations implies that this coordination mechanism may operate generally throughout much of the cortex and closely-associated structures. Co-ripples thus mark brief windows where increased firing within and between cortical and limbic locations could enable spike transmission between distant regions with sufficient temporal precision to drive plasticity (via spike timing dependent plasticity [40]) and information transfer via coincident firing [41] and related mechanisms [42–44].

We also note that co-ripple periods are very brief (∼100 ms), and the transient increase in firing rate during these windows may itself contribute to their functional role. Coordinated elevations in excitability across regions — even setting aside fine-timescale synchrony — could facilitate the inter-regional communication necessary for memory reinstatement. However, our STTC analyses demonstrate that synchrony during co-ripples is enhanced beyond what would be expected from rate increases alone, indicating that ripples coordinate the precise timing of spikes across regions rather than simply elevating excitability.

The specificity of these coordination effects to the ripple frequency band is also noteworthy. Our comparative analysis across frequency bands revealed that ripple oscillations (70-100 Hz) exhibited stronger enhancement of long-range coordination than oscillations in either low gamma (30-55 Hz) or non-oscillatory activity in very high gamma (120-190 Hz) ranges. Recently, across widespread cortical areas, ripple band co-oscillations were found to show much stronger modulation by a semantic judgement task than those in the very high gamma range [25]. This frequency specificity suggests that ripples occupy an optimal range for long-distance communication, potentially balancing the time needed for effective local processing with that required for propagation between co-rippling sites.

Previous work has shown that co-ripples are characterized by zero-lag phase-locking across large cortical areas during a reading task [25] and spontaneous waking [10]. They thus can be described as coupled oscillators, a common phenomenon in biology and physics [45]. Coupled oscillators are most often modeled with zero transmission delay between oscillating sites, whereas the distant locations in our recordings are connected by significant transmission delays. However, mathematical [46, 47] and computational neural [48] models confirm that coupled oscillators with significant delays can synchronize at zero-lag provided that multiple pathways between oscillators are present. Consistent with this framework, spiking neural network simulations of high-frequency bursts in the same spectral range have shown that both correlated external inputs and reciprocal connectivity are sufficient to produce synchronized activity across networks[5]. Other co-ripple characteristics have previously been reported that are consistent with coupled oscillators: the consistent center frequency especially at times of high phase-locking[25]; the exponential growth of co-rippling sites prior to the response[25]; and the lack of decrement with distance noted above. These findings suggest that the phenomena reported here result from activation of an interactive network rather than sequential directed transmission. Clearly, much experimental work remains to clarify the structure and dynamics of such a network.

Our findings also have implications for theories of cortical integration, which may be broadly classified as focal-sequential versus distributed-interactive, continuing the longstanding debate between localizationist and equipotentiality views of brain function [49]. While the focal-sequential framework has been successful in describing early sensory processing, such as the hierarchical organization of the ventral visual stream [50], it faces challenges in accounting for higher cognitive functions in association cortex. Approximately 75% of human cortex lies beyond the clearly hierarchical early sensory and late motor areas [51], and about 75% of reaction time in a simple semantic judgment task occurs after visual wordform encoding [25]. Many theories of cognitive processing posit a division of cortex into sequential-hierarchical and interactive-associative areas, with the later integrating various sensory, semantic, and executive information in guiding a response, and termed NeuroCognitive Networks [52, 53], Cognits [54], or the Global Neuronal Workspace [55]. A serious challenge to such theories has been the general lack of evidence for integration of neuronal firing over wide expanses of association cortex and closely-linked subcortical areas. Our study provides evidence for an essential component of that process, demonstrating task-related enhancement of specific neuronal firing between widely distributed cortical and subcortical structures in both hemispheres.

## Limitations

Importantly, electrode placement was determined entirely by clinical necessity, targeting regions most relevant for seizure monitoring rather than areas traditionally associated with working memory processing. Despite this constraint—which resulted in uneven regional coverage and prevented systematic testing of areas like posterior parietal cortex—we observed task-related modulation across all recorded sites. This widespread modulation, even in regions not classically implicated in working memory, supports theories of distributed cortical processing and suggests that cognitive functions engage broad neural networks rather than isolated specialized regions. While our sparse sampling likely captures only a subset of the broader coordination occurring during working memory, the consistent effects across diverse recording locations strengthen the evidence for ripples as a general mechanism of cortical communication.

Finally, our investigation was limited to the Sternberg working memory task with a narrow load manipulation (1 versus 3 items). Future studies employing diverse paradigms and broader task difficulty ranges would help establish whether ripple coordination represents a general cognitive mechanism or is specific to particular memory operations.

## Conclusion

In conclusion, our findings establish ripple oscillations as a mechanism for coordinating stimulus-specific neuronal firing across distant human brain regions during cognition. Co-ripples are not only associated with elevated excitability: they promote the reinstatement of encoding-specific co-firing patterns that predict behavioral efficiency, with effects that scale with cognitive demand and extend across brain-wide distances without significant attenuation. By providing direct single-neuron evidence for the spiking content carried during co-ripple windows, these results support a model in which co-ripple mediated coordination of unit firing serves as a fundamental mechanism for integrating distributed neural representations during human cognition.

## Methods

### Software

All post-processing and statistical analyses were performed using MATLAB 2019b (MathWorks, Natick, MA). Custom analysis code is available at https://github.com/iverzh/ripple-working-memory, and https://github.com/iverzh/ripple-detection.

### Participants and Data Collection

In this study, we analyzed data previously described by Daume et al. [27], which consists of intracranial recordings from 35 patients (43 sessions; 21 female, 14 male; age range 20-67 years) implanted with depth electrodes with embedded microwires for evaluation of medically refractory epilepsy. All patients and sessions from the original dataset were included in our study except for one patient (P88T), which was removed due to a lack of clean LFP microwire data in any of the implanted sites. Each patient was implanted with up to ten electrodes, comprised of a clinical macro-electrode with 8 microwires protruding ∼2-5 mm from the tip [56]. Electrodes considered for analysis targeted the ventromedial prefrontal cortex (vmPFC), anterior cingulate cortex (ACC), pre-supplementary motor area (preSMA), amygdala (AMY), and hippocampus (HIP), bilaterally.

Local field potentials (LFPs) and single-unit activity were simultaneously recorded at 32 kHz (ATLAS system, Neuralynx; Cedars-Sinai Medical Center and Toronto Western Hospital) or 30 kHz (Blackrock Neurotech; Johns Hopkins Hospital) from the implanted microwires. All recordings were locally referenced within each recording site using either one of the available microwire channels or a dedicated reference channel with lower impedance. The dataset included a total of 2253 microwire channels from which LFPs were recorded. All patients provided informed consent, and procedures were approved by the Institutional Review Board at each participating institution.

### Sternberg Memory Task

The task analyzed has been described in detail previously [27]. Briefly, patients performed a modified Sternberg working memory task with varying memory loads (1 or 3 items). In each trial (see Fig. 1c), they were instructed to remember a set of images presented sequentially (encoding phase), maintain the memory over a delay period (maintenance phase), and identify whether a retrieval image was part of the original set (retrieval phase). Each session consists of 140 trials.

The 280 images were chosen from five semantic categories (people, animals, landscapes, fruits, and either cars or tools depending on the version). Each trial sequence began with a variable-duration baseline point (900-1200ms), followed by a stimulus presentation phase. In the low-load condition (70 trials), participants viewed a single image for 2 seconds. In the high-load condition (70 trials), participants viewed a sequence of three images from different categories, each displayed for 2 seconds with brief interstimulus intervals ranging from 17-200ms. Following stimulus presentation, a retention interval commenced, signaled by the word “HOLD” appearing centrally for 2.5-2.8 seconds. This maintenance period was identical for both load conditions.

The trial concluded with a memory probe showing a single image, during which participants indicated whether the retrieval image matched one of the images shown in the current trial’s encoding phase. To prevent strategies based on image familiarity, non-matching retrieval images were always selected from earlier trials rather than introducing entirely novel images. These non-matching retrieval images were deliberately chosen from categories not represented in the current trial’s encoding set. Responses were collected using a Cedrus response pad, with response mapping reversed midway through the session following a brief rest period. For participants completing multiple sessions, entirely new image sets were used.

### Data Curation and Quality Control

To ensure high data quality and reliable analyses, we implemented a rigorous data curation protocol. All recordings were manually reviewed to identify and exclude channels contaminated with epileptiform activity, electrical artifacts, or excessive noise. For each patient, recordings were excluded from time periods containing interictal epileptiform discharges spreading across multiple channels, or periods of increased noise due to patient movement. Electrodes substantially contaminated by epileptiform activity were not used for analysis.

Artifact and interictal discharge detection was performed on a per-wire basis using a semiautomated algorithm together with subsequent visual inspection. To detect high-amplitude noise events and inter-ictal discharges, we z-scored the amplitude in each channel across all trials. To avoid amplitude biasing from extreme values, we first capped the data at 6 standard deviations from the mean and then re-performed z-scoring on the capped data. Trials in which any time sample exceeded a threshold of 4 standard deviations were removed from the analysis for that wire. Signal jumps were detected by z-scoring the difference between every fourth sample of the capped signal, with trials containing jumps exceeding 10 standard deviations being excluded.

### Electrodes and Visualization

Electrode locations were used as stated in the dataset (Fig 1.A). Electrode locations were distributed across frontal and temporal regions, allowing for the analysis of both local and long-range interactions. We estimated cross bundle fiber tract distances according to probabilistic diffusion MRI white matter tractography [57], computed from population averages [58]. To determine the appropriate streamline lengths, each bundle was associated with a parcel from the HCP-MMP1.0 atlas [59]. Specifically, bundles implanted in the vmPFC, ACC, preSMA, AMY and HIP sites were assigned to the s32, p24, 8BM, TGd and H parcels, respectively.

### Detection of Ripples and Oscillations in Other Gamma Ranges

Ripple detection was performed on local field potential data from each microwire electrode, downsampled to 1000 Hz, and using established methods adapted from previous studies [25, 26]. Data were bandpass filtered using a sixth-order Butterworth filter in the ripple frequency range (70-100 Hz, applied forward and reverse for zero-phase distortion), then z-scored relative to the entire recording session. Initial candidate events were identified as containing at least three consecutive ripple cycles with z-scores exceeding 1.

Candidate events underwent further selection criteria, requiring the maximum z-score of the analytic amplitude in the ripple band to exceed 2.5 standard deviations. Ripple events occurring within 25 ms of each other were merged into single events. Ripple centers were defined as the timepoint of maximum positive deflection in the ripple-band filtered signal.

For determining event boundaries, the z-scored ripple-band analytic amplitude was smoothed using a 100 ms sliding Gaussian kernel. Event onsets and offsets were marked when this smoothed amplitude envelope dropped below a z-score threshold of 0.75. Ripples were excluded if the 100-Hz high-pass z-score exceeded 7 in absolute value or if they occurred within 2 seconds of large voltage deflections (≥3 mV/ms). Events were also rejected if they fell within ±500 ms of detected interictal spikes [11, 26].

Additional quality control measures included excluding events with single prominent cycles or those where the largest valley-to-peak amplitude in the broadband signal exceeded 2.5 times the third-largest amplitude. For each recording channel, the average ripple-triggered LFP was visually inspected to verify multiple distinct cycles at ripple frequency, and time-frequency spectrograms were examined to confirm discrete power increases within the ripple band. Individual ripple events and co-occurring ripples across channels were manually reviewed to ensure multiple ripple-frequency cycles without contamination from artifacts, unit activity bleed-through, or epileptiform discharges.

For comparison, we also detected oscillatory events in low gamma (30-55 Hz) and very high gamma (120-190 Hz) frequency bands using the same thresholding methodology, adjusting the filter bands accordingly. We characterized the reliability of these signals to differentiate between memory load conditions using the contrast-to-noise ratio (CNR):

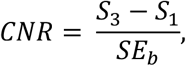

where *S_3_* and *S_1_* are the oscillation occurrence rates in load 3 and load 1 respectively, and *Se_b_* is the standard error across all subjects and trials during the 1s baseline period.

### Time Frequency Analysis

Time-frequency analysis of ripple-associated spectral dynamics was performed using broadband local field potential recordings processed through EEGLAB [60]. We computed event-related spectral perturbations (ERSP) across frequencies ranging from 1 to 500 Hz at 1 Hz intervals, with ripple onset aligned to time zero. The analysis employed windowed fast Fourier transforms using Hanning tapers, which were subsequently averaged to generate mean time-frequency representations.

Baseline normalization was applied to each frequency bin by dividing spectral power values by the average power recorded during the pre-event period (−2000 to −1500 ms). Statistical significance testing utilized two-tailed bootstrap procedures (200 iterations) with multiple comparison correction via false discovery rate (FDR) control at α=0.05.

### Detection and Selection of Single Units

Isolated single neurons were used as provided in the dataset [27]. We applied additional exclusion criteria:

Units were excluded if the mean and distribution of spike waveforms exhibited low signal-to-noise characteristics, defined as a peak signal-to-noise ratio of 2.5 or less. Additionally, we excluded units whose auto-correlograms indicated contamination from multiple neurons, specifically those with greater than 3% of spikes occurring within 3 ms of each other. This 3 ms criterion corresponds to the typical refractory period of neuronal firing, during which a single neuron cannot generate consecutive action potentials. Units violating this criterion likely represent multi-unit activity, which could confound analyses of precise spike timing relationships essential for co-firing analyses. These conservative selection criteria ensured that our dataset comprised well-isolated single neurons with reliable spike timing, critical for examining the temporal relationships between unit activity and ripple oscillations.

The flexible positioning of microwires within each electrode bundle can occasionally result in multiple wires detecting signals from the same neuron, especially when slight wire movements bring them into closer proximity. To prevent this from confounding our analyses, we systematically identified and merged units across different wires within the same bundle that likely originated from a single neuronal source. Units exhibiting greater than 30% of spikes occurring within 1 ms of each other were merged, as this exceptionally high temporal correlation indicates recording from the same neuron rather than genuine co-firing between distinct cells. This merging procedure was essential for preventing artificial inflation of co-firing rates and ensuring that all observed neural coordination reflected authentic interactions between separate neuronal units.

Following these quality control procedures, we successfully isolated 1,373 high-quality single units distributed across all recorded brain regions: 360 units in the hippocampus, 496 in the amygdala, 206 in the ventromedial prefrontal cortex, 188 in the dorsal anterior cingulate cortex, and 204 in the pre-supplementary motor area.

### Spike Removal from LFP

While local field potentials reflect slower network oscillations, the high-amplitude nature of action potentials can introduce significant artifacts into LFP recordings, particularly in higher frequency bands where spike-LFP coupling analyses are performed. This contamination occurs because the large voltage deflections associated with individual spikes can influence the filtered LFP signal across a broad frequency spectrum, creating spurious correlations between unit activity and oscillatory patterns [61].

To address this potential confound, we implemented a spike removal procedure prior to LFP processing. For each identified unit, we calculated the average spike waveform and subtracted this template from the raw 30 kHz signal at every detected spike time from – 1ms to + 2ms around a the peak of the spike. This preprocessing step was applied before down-sampling and frequency filtering operations, ensuring that subsequent LFP analyses reflected genuine network oscillations rather than artifacts from unit activity. This approach preserves the integrity of ripple-unit relationships while eliminating the influence of spike waveform contamination on oscillatory measurements, allowing for more accurate assessment of true physiological relationships between neural firing and local field oscillations.

Even after removal of detected spike waveforms, spikes from more distant cells could summate and contribute to the ripple band signal. A modeling study of rat hippocampus that included adjacent as well as distant action-potentials calculated that about half of the ripple amplitude is due to synaptic currents and half to action-potentials at 160Hz, the frequency of rodent ripples [62]. However, the synaptic proportion was greater at 90Hz (the frequency of human ripples). Furthermore, there are many critical parameters in their model that are very different in rodents vs humans, and in hippocampus vs cortex or amygdala: pyramidal cell size and morphology, firing rates during ripples (human PY fire less than once per ripple), packing density, connectivity, synaptic and cellular time-constants, etc. In the absence of modeling appropriate to the species and sites reported here, it appears likely that the majority of the ripple signal is due to synaptic currents. More generally, unit firing is phase-modulated by ripple oscillations, and the two co-occur; the exact contribution of each to the recorded ripple-band signal depends on multiple biophysical factors — including cell morphology, firing rates, and local circuit properties — as well as electrode size, shape, and location. Units and LFP thus provide distinct measures of activity during ripples, consistent with the overall correlation of firing with ripples reported above indicating that ripple rates during maintenance and retrieval explain only ∼5% of the variance in firing rate in these periods.

### Co-ripple Rates and Co-firing Calculation

Co-occurring ripples (co-ripples) were defined as ripple events that temporally overlapped by at least 25ms across two different microwire bundles. We quantified co-ripple rates as the percentage of ripples on one electrode that co-occurred with ripples on another electrode. We examined ripple co-occurrence within microwire bundles (intra-bundle, <∼5 mm separation) and across different microwire bundles within the same hemisphere (inter-bundle, 71-203 mm separation) and across hemispheres (35-223 mm separation).

To measure neural co-firing between distant brain regions, we defined co-firing as the occurrence of spikes from units in separate brain regions within a 25ms window. Co-ripple periods were defined as times when ripples were simultaneously detected on microwires in both units’ respective bundles. Conversely, no-ripple periods were defined as times when no ripples were detected in either bundle. For across-bundle unit pairs, we categorized them into five connection types: amygdala-cortex (AMY-CTX), hippocampus-cortex (HIP-CTX), amygdala-hippocampus (AMY-HIP), ipsilateral cortico-cortical (CTX ipsi), and contralateral cortico-cortical (CTX contra).

For each cell-pair, we calculated the co-firing rate during co-ripple periods and no-ripple periods by counting the number of co-occurring spikes and dividing by the total duration of the respective periods. To account for potential bias from differences in baseline firing rates across brain regions, we also computed normalized co-firing rates by dividing the observed co-firing rate by the mean of the individual firing rates of each unit pair (Supplementary Fig. 2).

### Within-Bundle and Across-Bundle Ripple Response Curves

To examine how ripple activity varies with cognitive demand and behavioral performance, we computed ripple overlap rates both within individual microwire bundles and across different bundles under various task conditions.

For within-bundle ripple rate analysis (Fig. 2), we calculated the ripple density as the number of ripples across all microwires within each bundle that overlapped in 1 millisecond bins and averaged across all trials and participants, yielding the mean total ripples occurring within each bundle per trial (ie., ripples trial^-1^). These response curves were computed separately for high memory load (3 items) and low memory load (1 item) trials. To assess the relationship between ripple activity and behavioral performance, we further subdivided load 3 trials based on response time. For each session, we calculated the median response time for all load 3 trials, then classified trials as “fast” (below median) or “slow” (above median). Unlike memory load conditions which were experimentally balanced, fast and slow response trials varied naturally across participants, and each participant had different numbers of useable microwires per bundle. Therefore, when comparing response time effects, we normalized ripple rates by the number of viable microwires per bundle (ie., ripples trial^-1^ wire^-1^) to prevent bias from unequal sampling across participants.

For across-bundle co-ripple analysis by memory load (Fig. 3), we identified periods of the recording where at least one ripple was simultaneously detected across two different microwire bundles. In contrast to the within-bundle analysis, any periods where only one bundle was rippling were considered as non-co-rippling periods. To quantify co-ripple activity, we counted the number of microwires in each bundle that contained overlapping ripples, binned this count in 1 ms windows, and then averaged across all trials and participants to obtain a co-ripple rate (co-ripples trial^-1^). This metric was computed separately for each bundle pair and memory load condition to assess how cognitive demand modulates inter-regional ripple coordination.

### Rate-Independent Measures of Pairwise Neural Synchrony

To assess whether enhanced co-firing during co-ripple events reflects true increases in neural synchrony beyond what would be expected from elevated firing rates alone, we employed two complementary approaches: the spike time tiling coefficient (STTC) and a simpler analytic null model comparison.

The STTC quantifies pairwise correlations between spike trains while controlling for differences in firing rate between neurons and across conditions [29]. For two cells A and B, the coefficient is calculated as:

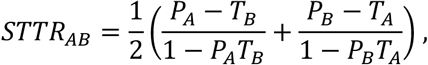

where P_A_ represents the proportion of spikes from cell A falling within ±Δt of any spike from cell B (and vice versa for P_B_), and T_A_ represents the proportion of total recording time covered by ±Δt windows around cell A’s spikes (and vice versa for T_B_). We used Δt = 25 ms. STTC values are bounded between −1 and +1, with positive values indicating correlated firing, zero indicating independence, and negative values indicating anti-correlation.

Because STTC was originally validated on continuous recordings of at least 10 minutes duration [29] and our co-ripple periods averaged only approximately 45 seconds of total time per recording, we implemented inclusion criteria to ensure reliable estimation. Specifically, we restricted the STTC analysis to neuron pairs exhibiting at least one co-firing event (spikes within ±Δt) during both co-ripple and no-ripple periods. This criterion ensured that STTC values reflected meaningful spike timing relationships rather than undefined or unstable estimates arising from sparse data. STTC was computed separately for co-ripple and no-ripple periods across regional pairings: cortical-cortical (ctx-ctx), limbic-cortical (lim.-ctx), limbic-limbic (lim.-lim.), ipsilateral (ipsi), and contralateral (contra) cell pairs.

As a complementary approach that permitted inclusion of all recorded cell pairs regardless of co-firing occurrence, we computed the observed co-firing rate relative to an analytic null model. The null model estimates the expected co-firing rate (ECFR) under the assumption of independent Poisson firing:

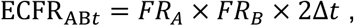

where FR_A_ and FR_B_ are the firing rates of cells A and B, respectively, and Δt defines the coincidence window (25 ms). We then computed the difference between the observed co-firing rate and this null expectation (co-fire rate above null model, reported in ΔHz). Positive values indicate co-firing in excess of that expected from firing rates alone, providing a firing rate-independent measure of synchrony. This analysis included all cell pairs, as the null model remains well-defined even in the absence of observed co-firing events.

### Calculation of Task Phase and Load Modulation of Co-firing

We examined how ripple occurrence and associated co-firing varied across different cognitive phases by measuring percentage changes relative to a pre-trial baseline period (spanning −0.9 to −0.3 seconds before the first stimulus appeared). This baseline comparison allowed us to identify which task phases—encoding, maintenance, or retrieval—showed the greatest enhancement of synchronized neural activity, thereby revealing how memory processes selectively recruit coordinated activity across distributed brain networks during distinct cognitive operations.

To assess how memory demands influence neural coordination between brain regions, we analyzed co-firing rates under different conditions: during co-ripple periods (when ripples occurred simultaneously in both regions), during no-ripple periods (when no ripples were detected in either region), and across all time periods combined. For each analysis, we compared co-firing between high memory load (load 3) and low memory load (load 1) trials.

For every unit pair, we calculated the percentage change in co-firing from load 1 to load 3 trials using the formula:

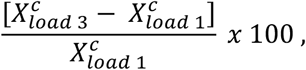

where *X* is the number of co-firing events during a given load and during the ripple condition *c* (i.e., co-ripple, no-ripple and all periods). To ensure statistical reliability, we restricted our analysis to unit pairs with a baseline co-firing rate of at least 0.025 Hz, yielding 12,335 analyzable pairs from the total pool of 16,274 recorded unit pairs.

### Repeated Stimulus Co-firing

To investigate whether co-ripples support the reinstatement of stimulus-specific neural representations during memory retrieval, we analyzed the repetition of co-firing patterns between encoding and retrieval phases. For each cell pair recorded from different brain regions, we identified co-firing events, defined as spikes occurring within a 25 ms temporal window between the two units.

We tracked co-firing patterns separately for each stimulus presented during the task. During the encoding phase (0 – 2 second post stimulus onset), we recorded which cell pairs exhibited co-firing when each specific stimulus was presented. During the retrieval phase (0 – 1 second post probe stimulus onset), we then examined whether these same cell pairs showed co-firing when either the same stimulus (match trials) or a different stimulus (mismatch trials) was presented.

The analysis calculated the percentage of cell-pair trials (# cell-pairs × # total trials) that exhibited repeated co-firing patterns between encoding and retrieval phases as:

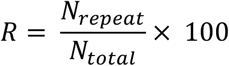

Where, *R* = repetition rate (percentage), *N_repeat_ =* number of cell-pair trials with co-firing at both encoding and retrieval phases, and *N_total_* = total number of cell pair trials. A cell-pair trial was defined as a unique combination of a specific cell pair during a given task trial (i.e., recording from 100 cell pairs across 50 trials, would yield 5,000 cell-pair trials). *R* was computed during load 3 trials separately for fast responses and slow response, defined as the top and bottom 50% of response times.

To determine the specific contribution of co-ripples to repetition of co-firing patterns, we separated our analysis into co-ripple periods and no-ripple periods. Co-ripple periods were defined as times when ripples were simultaneously detected in both brain regions containing the analyzed unit pair. No-ripple periods were duration-matched to co-ripple periods through 25 random shuffles per cell-pair trial to ensure equivalent temporal sampling.

### Stimulus-label shuffle control for co-firing reinstatement analysis

To assess whether co-firing reinstatement during co-ripples reflected stimulus-specific pattern matching rather than generally elevated excitability, we performed a within-subject, trial-shuffled permutation test. For each cell pair, we identified all co-firing events (spikes within 25 ms) occurring during co-ripple periods in both the encoding and retrieval phases. We then shuffled the stimulus labels associated with encoding-period co-firing events, breaking the correspondence between specific stimuli and their associated co-firing patterns while preserving the overall co-firing rate observed during co-ripples. For each permutation, we recalculated the percentage of cell-pair-trials showing repeated co-firing between encoding and retrieval for the (now mismatched) stimulus labels. This procedure was repeated 1000 times to generate a null distribution of expected pattern match rates. Because the shuffled baseline is derived from the same co-ripple periods as the observed data, any above-chance pattern reinstatement cannot be attributed to differences in overall excitability between co-ripple and no-ripple periods. Significance was assessed by comparing the observed reinstatement rate to the 95% confidence interval of the shuffled null distribution.

### Temporal Relationship Between Co-rippling and Repeated co-firing

To examine the temporal relationship between stimulus-specific co-firing and co-ripples, we constructed peri-ripple time histograms to visualize when matching co-fire events occurred relative to co-ripple timing. This temporal analysis was conducted separately for unit pairs within the same hemisphere and those across different hemispheres, allowing us to assess whether the temporal coupling of stimulus-specific firing differed based on anatomical distance.

We first identified cell pairs that co-fired during the encoding phase for each stimulus. We then identified all instances where those same cell pairs co-fired again during the retrieval phase when that stimulus was probed—constituting stimulus-specific reinstatement events. Critically, these matching co-fire events were not selected based on their timing relative to ripples and could occur at any point during the probe period.

For each detected co-ripple (defined as ripples occurring simultaneously in both recorded regions), we identified the temporal center of the overlapping period and created a time window extending ±200 ms around this center. We then counted all instances where the same cell pairs that co-fired during encoding of a specific stimulus also co-fired during the retrieval phase presentation of that same stimulus, binning these events by their temporal offset from the co-ripple center. This analysis tests whether stimulus-specific reinstatement events are temporally clustered around co-ripples, which is not guaranteed by the selection criteria.

To establish the specificity of this temporal relationship, we implemented two control analyses. First, we repeated the same procedure for single ripples that occurred in only one of the two bundle locations while no ripple was detected in the other location, allowing us to determine whether the temporal coupling required simultaneous rippling in both regions. Second, we created surrogate data by randomly shuffling spike times within each trial while preserving the overall firing rate profiles and inter-spike interval distributions. Importantly, spike times were shuffled separately within the encoding and retrieval stages for each trial, maintaining the temporal structure within each task phase. We then recomputed the enrichment of matching co-fire events around the original co-ripple centers using these shuffled spike trains.

### Statistical Methods

All statistical analyses were performed on data from 35 patients across 43 recording sessions during a modified Sternberg working memory task. Single units were identified and classified across 1927 microwire channels, yielding 1373 isolated units. For analyses examining unit co-firing between brain regions, we included only unit pairs with a baseline co-firing rate of at least 0.025 Hz, resulting in 12,335 out of 16,274 total cell pairs being analyzed.

To assess task-related modulation of ripple rates and unit firing, we used permutation tests of the median to compare activity during encoding, maintenance, and retrieval phases against pre-trial baseline levels. The relationship between ripple rates and neuronal firing rates was evaluated using Pearson correlation coefficients across all task phases. For comparing co-firing rates between co-ripple and no-ripple periods across all unit pairs (n = 31,489), we employed paired permutation tests. Differences in co-firing between anatomical connection types (amygdala-cortex, hippocampus-cortex, amygdala-hippocampus, ipsilateral cortico-cortical, and contralateral cortico-cortical) were also assessed using paired permutation tests.

Memory load effects on ripple occurrence/co-occurrence and unit activity were analyzed using linear mixed-effects models with FDR correction for multiple comparisons. We examined how each neural signal was load-modulated by modelling the load condition as a fixed effect and the patient as a random effect according to the following:

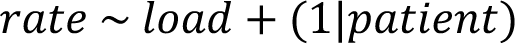

These models tested whether ripple rates, co-ripple rates, and firing rates differed between high memory load (3 items) and low memory load (1 item) conditions during maintenance and retrieval phases. The same approach was used to identify region pairs showing significant load-dependent modulation of oscillatory co-occurrence across different frequency bands (ripples, low gamma, and very high gamma).

The same model was used to test for significance in the relationship between ripple occurrence during the retrieval stimulus and response timing. Instead of using memory load, we categorized each trial as fast RT or slow RT (bottom and top 50% of response times, respectively).

For analyses of unit co-firing modulation by task phase and memory load, we used paired permutation tests to compare median co-firing rates during different task stages relative to baseline. For all permutation-based testing, 10000 iterations were performed. Kruskal-Wallis tests were employed to assess overall differences between co-ripple, no-ripple, and all-period conditions across task stages. Percent change in load-dependent co-firing were evaluated using paired one-sided Student’s t-tests comparing high versus low memory load conditions.

The analysis of stimulus-specific co-firing pattern repetition utilized chi-squared tests to compare the proportion of cell-pair trials exhibiting repeated co-firing patterns between encoding and retrieval phases for match trials (same stimulus) versus mismatch trials (different stimulus). This analysis was performed separately for co-ripple and no-ripple periods, with no-ripple periods duration-matched across 25 shuffles per cell-pair trial.

The statistical significance of co-firing enrichment around ripple events was assessed using one-way ANOVA across the three conditions (co-ripples, single ripples, shuffled controls). This analysis was performed separately for within-hemisphere and across-hemisphere connections during both encoding and retrieval stages, allowing us to determine whether co-ripples provide privileged temporal windows for reinstating stimulus-specific firing patterns across different anatomical pathways and task phases.

## Acknowledgements

We thank Sierra Wilson, Adam Niese, and Jacob Garrett for their support. This work was supported by NIMH (T32 MH020002, F31 MH135645). Acquisition, processing, and public release of the data was supported by the NIH BRAIN initiative through U01NS117839.

## Notes

https://dandiarchive.org/dandiset/000673

### Competing Interest Statement

The authors have declared no competing interest.

### Summary of Updates

first round of paper revisions. New analyses, figures and discussion.

## References

1. Fries P. Neuronal gamma-band synchronization as a fundamental process in cortical computation. Annu Rev Neurosci. 2009;32:209–24. doi: 10.1146/annurev.neuro.051508.135603.

2. Uhlhaas PJ, Pipa G, Lima B, Melloni L, Neuenschwander S, Nikolić D, et al. Neural synchrony in cortical networks: history, concept and current status. Front Integr Neurosci. 2009;3:17. doi: 10.3389/neuro.07.017.2009. PubMed Central PMCID: PMCPMC2723047.

3. Singer W. Neuronal synchrony: a versatile code for the definition of relations? Neuron. 1999;24(1):49–65, 111-25. doi: 10.1016/s0896-6273(00)80821-1.

4. Palmigiano A, Geisel T, Wolf F, Battaglia D. Flexible information routing by transient synchrony. Nat Neurosci. 2017;20(7):1014-+. doi: 10.1038/nn.4569. PubMed PMID: WOS:000404115100019.

5. Banaie Boroujeni K, Helfrich RF, Fiebelkorn IC, Bentley JN, Brunner P, Lin JJ, et al. High-frequency bursts facilitate fast communication for human spatial attention. Nat Neurosci. 2026;29(2):435–44. Epub 20251202. doi: 10.1038/s41593-025-02160-5. PubMed PMID: 41331141; PubMed Central PMCID: PMCPMC12695012.

6. Kucewicz MT, Cimbalnik J, Garcia-Salinas JS, Brazdil M, Worrell GA. High frequency oscillations in human memory and cognition: a neurophysiological substrate of engrams? Brain. 2024;147(9):2966–82. doi: 10.1093/brain/awae159. PubMed Central PMCID: PMCPMC11370809.

7. Roelfsema PR. Solving the binding problem: Assemblies form when neurons enhance their firing rate-they don’t need to oscillate or synchronize. Neuron. 2023;111(7):1003–19. doi: 10.1016/j.neuron.2023.03.016.

8. Buzsáki G. Hippocampal sharp wave-ripple: A cognitive biomarker for episodic memory and planning. Hippocampus. 2015;25(10):1073–188. doi: 10.1002/hipo.22488. PubMed Central PMCID: PMCPMC4648295.

9. Girardeau G, Benchenane K, Wiener SI, Buzsáki G, Zugaro MB. Selective suppression of hippocampal ripples impairs spatial memory. Nat Neurosci. 2009;12(10):1222–3. doi: 10.1038/nn.2384.

10. Dickey CW, Verzhbinsky IA, Jiang X, Rosen BQ, Kajfez S, Eskandar EN, et al. Cortical Ripples during NREM Sleep and Waking in Humans. J Neurosci. 2022;42(42):7931–46. doi: 10.1523/JNEUROSCI.0742-22.2022. PubMed Central PMCID: PMCPMC9617618.

11. Dickey CW, Verzhbinsky IA, Jiang X, Rosen BQ, Kajfez S, Stedelin B, et al. Widespread ripples synchronize human cortical activity during sleep, waking, and memory recall. Proc Natl Acad Sci U S A. 2022;119(28):e2107797119. doi: 10.1073/pnas.2107797119. PubMed Central PMCID: PMCPMC9282280.

12. Dickey CW, Verzhbinsky IA, Kajfez S, Rosen BQ, Gonzalez CE, Chauvel PY, et al. Thalamic spindles and Up states coordinate cortical and hippocampal co-ripples in humans. PLoS Biol. 2024;22(11):e3002855. doi: 10.1371/journal.pbio.3002855. PubMed Central PMCID: PMCPMC11575773.

13. Vaz AP, Inati SK, Brunel N, Zaghloul KA. Coupled ripple oscillations between the medial temporal lobe and neocortex retrieve human memory. Science. 2019;363(6430):975–8. doi: 10.1126/science.aau8956. PubMed Central PMCID: PMCPMC6478623.

14. Mishra A, Tostaeva G, Nentwich M, Espinal E, Markowitz N, Winfield J, et al. Motifs of human high-frequency oscillations structure processing and memory of continuous audiovisual narratives. Science Advances. 2025;11(30). doi: ARTN eadv0986 10.1126/sciadv.adv0986. PubMed PMID: WOS:001535436800005.

15. Xie W, Wittig JH, Chapeton JI, El-Kalliny M, Jackson SN, Inati SK, et al. Neuronal sequences in population bursts encode information in human cortex. Nature. 2024:1–8. doi: 10.1038/s41586-024-08075-8.

16. Wilson MA, McNaughton BL. Reactivation of hippocampal ensemble memories during sleep. Science. 1994;265(5172):676–9. doi: 10.1126/science.8036517.

17. Foster DJ, Wilson MA. Reverse replay of behavioural sequences in hippocampal place cells during the awake state. Nature. 2006;440(7084):680–3. doi: 10.1038/nature04587.

18. Nitzan N, McKenzie S, Beed P, English DF, Oldani S, Tukker JJ, et al. Propagation of hippocampal ripples to the neocortex by way of a subiculum-retrosplenial pathway. Nat Commun. 2020;11(1):1947. doi: 10.1038/s41467-020-15787-8. PubMed Central PMCID: PMCPMC7181800.

19. Nitzan N, Swanson R, Schmitz D, Buzsáki G. Brain-wide interactions during hippocampal sharp wave ripples. Proc Natl Acad Sci U S A. 2022;119(20):e2200931119. doi: 10.1073/pnas.2200931119. PubMed Central PMCID: PMCPMC9171920.

20. Khodagholy D, Gelinas JN, Buzsáki G. Learning-enhanced coupling between ripple oscillations in association cortices and hippocampus. Science. 2017;358(6361):369–72. doi: 10.1126/science.aan6203. PubMed Central PMCID: PMCPMC5872145.

21. Doostmohammadi J, Gieselmann MA, van Kempen J, Lashgari R, Yoonessi A, Thiele A. Ripples in macaque V1 and V4 are modulated by top-down visual attention. Proc Natl Acad Sci U S A. 2023;120(5):e2210698120. doi: 10.1073/pnas.2210698120. PubMed Central PMCID: PMCPMC9945997.

22. Vaz AP, Wittig JH, Jr., Inati SK, Zaghloul KA. Replay of cortical spiking sequences during human memory retrieval. Science. 2020;367(6482):1131–4. doi: 10.1126/science.aba0672. PubMed Central PMCID: PMCPMC7211396.

23. Tong APS, Vaz AP, Wittig JH, Inati SK, Zaghloul KA. Ripples reflect a spectrum of synchronous spiking activity in human anterior temporal lobe. Elife. 2021;10. doi: 10.7554/eLife.68401. PubMed Central PMCID: PMCPMC8716101.

24. Norman Y, Yeagle EM, Khuvis S, Harel M, Mehta AD, Malach R. Hippocampal sharp-wave ripples linked to visual episodic recollection in humans. Science. 2019;365(6454). doi: 10.1126/science.aax1030.

25. Garrett JC, Verzhbinsky IA, Kaestner E, Carlson C, Doyle WK, Devinsky O, et al. Binding of cortical functional modules by synchronous high-frequency oscillations. Nature Human Behaviour. 2024:1–15. doi: 10.1038/s41562-024-01952-2.

26. Verzhbinsky IA, Rubin DB, Kajfez S, Bu Y, Kelemen JN, Kapitonava A, et al. Co-occurring ripple oscillations facilitate neuronal interactions between cortical locations in humans. Proc Natl Acad Sci U S A. 2024;121(1):e2312204121. doi: 10.1073/pnas.2312204121.

27. Daume J, Kamiński J, Schjetnan AGP, Salimpour Y, Khan U, Kyzar M, et al. Control of working memory by phase-amplitude coupling of human hippocampal neurons. Nature. 2024;629(8011):393–401. doi: 10.1038/s41586-024-07309-z. PubMed Central PMCID: PMC2925295.

28. Dickey CW, Sargsyan A, Madsen JR, Eskandar EN, Cash SS, Halgren E. Travelling spindles create necessary conditions for spike-timing-dependent plasticity in humans. Nat Commun. 2021;12(1):1–15. doi: 10.1038/s41467-021-21298-x.

29. Cutts CS, Eglen SJ. Detecting pairwise correlations in spike trains: an objective comparison of methods and application to the study of retinal waves. J Neurosci. 2014;34(43):14288–303. doi: 10.1523/JNEUROSCI.2767-14.2014. PubMed PMID: 25339742; PubMed Central PMCID: PMCPMC4205553.

30. Kunz L, Staresina BP, Reinacher PC, Brandt A, Guth TA, Schulze-Bonhage A, et al. Ripple-locked coactivity of stimulus-specific neurons and human associative memory. Nat Neurosci. 2024. doi: 10.1038/s41593-023-01550-x. PubMed Central PMCID: PMC4194129.

31. Sternberg S. In defence of high-speed memory scanning. Q J Exp Psychol (Hove). 2016;69(10):2020–75. doi: 10.1080/17470218.2016.1198820. PubMed PMID: 27557823.

32. Townsend JT, Fific M. Parallel versus serial processing and individual differences in high-speed search in human memory. Percept Psychophys. 2004;66(6):953–62. doi: 10.3758/bf03194987. PubMed PMID: 15675643.

33. McClelland JL, McNaughton BL, Lampinen AK. Integration of new information in memory: new insights from a complementary learning systems perspective. Philos T R Soc B. 2020;375(1799). doi: ARTN 20190637 10.1098/rstb.2019.0637. PubMed PMID: WOS:000523698100013.

34. Yonelinas A, Hawkins C, Abovian A, Aly M. The role of recollection, familiarity, and the hippocampus in episodic and working memory. Neuropsychologia. 2024;193. doi: ARTN 108777 10.1016/j.neuropsychologia.2023.108777. PubMed PMID: WOS:001166171100001.

35. Lundqvist M, Herman P, Warden MR, Brincat SL, Miller EK. Gamma and beta bursts during working memory readout suggest roles in its volitional control. Nat Commun. 2018;9(1):394. doi: 10.1038/s41467-017-02791-8. PubMed Central PMCID: PMCPMC5785952.

36. Daume J, Kamiński J, Salimpour Y, Gómez Palacio Schjetnan A, Anderson WS, Valiante TA, et al. Persistent activity during working memory maintenance predicts long-term memory formation in the human hippocampus. Neuron. 2024;0(0). doi: 10.1016/j.neuron.2024.09.013.

37. Donahue CJ, Sotiropoulos SN, Jbabdi S, Hernandez-Fernandez M, Behrens TE, Dyrby TB, et al. Using Diffusion Tractography to Predict Cortical Connection Strength and Distance: A Quantitative Comparison with Tracers in the Monkey. J Neurosci. 2016;36(25):6758–70. doi: 10.1523/JNEUROSCI.0493-16.2016. PubMed PMID: 27335406; PubMed Central PMCID: PMCPMC4916250.

38. Rosen BQ, Halgren E. An estimation of the absolute number of axons indicates that human cortical areas are sparsely connected. PLoS Biol. 2022;20(3):e3001575. Epub 20220314. doi: 10.1371/journal.pbio.3001575. PubMed PMID: 35286306; PubMed Central PMCID: PMCPMC8947121.

39. Buzsáki G, Draguhn A. Neuronal oscillations in cortical networks. Science. 2004;304(5679):1926–9. doi: DOI 10.1126/science.1099745. PubMed PMID: WOS:000222241600031.

40. Feldman DE. The spike-timing dependence of plasticity. Neuron. 2012;75(4):556–71. doi: 10.1016/j.neuron.2012.08.001. PubMed Central PMCID: PMCPMC3431193.

41. Salinas E, Sejnowski TJ. Correlated neuronal activity and the flow of neural information. Nat Rev Neurosci. 2001;2(8):539–50. doi: Doi 10.1038/35086012. PubMed PMID: WOS:000170410300012.

42. Panzeri S, Moroni M, Safaai H, Harvey CD. The structures and functions of correlations in neural population codes. Nat Rev Neurosci. 2022;23(9):551–67. doi: 10.1038/s41583-022-00606-4. PubMed PMID: WOS:000814474600002.

43. Dahmen D, Layer M, Deutz L, Dabrowska PA, Voges N, von Papen M, et al. Global organization of neuronal activity only requires unstructured local connectivity. Elife. 2022;11. doi: ARTN e68422 10.7554/eLife.68422. PubMed PMID: WOS:000794921600001.

44. Schneidman E, Berry MJ, Segev R, Bialek W. Weak pairwise correlations imply strongly correlated network states in a neural population. Nature. 2006;440(7087):1007–12. doi: 10.1038/nature04701. PubMed PMID: WOS:000236906000026.

45. Ermentrout B, Park Y, Wilson D. Recent advances in coupled oscillator theory. Philos Trans A Math Phys Eng Sci. 2019;377(2160):20190092. Epub 20191028. doi: 10.1098/rsta.2019.0092. PubMed PMID: 31656142; PubMed Central PMCID: PMCPMC7032540.

46. Esfahani ZG, Gollo LL, Valizadeh A. Stimulus-dependent synchronization in delayed-coupled neuronal networks. Scientific Reports. 2016;6(1):23471. doi: 10.1038/srep23471.

47. Ghadiri A, Rezaei H, Aertsen A, Kumar A, Valizadeh A. Delay selection by spike-timing-dependent plasticity shapes efficient networks for signal transmission. bioRxiv. 2025:2025.10.20.682868. doi: 10.1101/2025.10.20.682868.

48. Bazhenov M, Rulkov NF, Timofeev I. Effect of synaptic connectivity on long-range synchronization of fast cortical oscillations. J Neurophysiol. 2008. PubMed PMID: 18632897.

49. García-Molina A, Peña-Casanova J. Functional organisation of the cerebral cortex: from Gall to Lashley. 2024.

50. DiCarlo JJ, Zoccolan D, Rust NC. How Does the Brain Solve Visual Object Recognition? Neuron. 2012;73(3):415–34. doi: 10.1016/j.neuron.2012.01.010. PubMed PMID: WOS:000300140600005.

51. Hayashi T, Hou Y, Glasser MF, Autio JA, Knoblauch K, Inoue-Murayama M, et al. The nonhuman primate neuroimaging and neuroanatomy project. Neuroimage. 2021;229:117726. doi: 10.1016/j.neuroimage.2021.117726. PubMed Central PMCID: PMCPMC8079967.

52. Mesulam M. Representation, Inference, and Transcendent Encoding in Neurocognitive Networks of the Human Brain. Ann Neurol. 2008;64(4):367–78. doi: 10.1002/ana.21534. PubMed PMID: WOS:000260845000005.

53. Bressler SL, Richter CG. Interareal oscillatory synchronization in top-down neocortical processing. Curr Opin Neurobiol. 2015;31:62–6. Epub 20140915. doi: 10.1016/j.conb.2014.08.010. PubMed PMID: 25217807.

54. Fuster JM. Cognitive Networks (Cognits) Process and Maintain Working Memory. Front Neural Circuits. 2021;15:790691. Epub 20220118. doi: 10.3389/fncir.2021.790691. PubMed PMID: 35115910; PubMed Central PMCID: PMCPMC8803648.

55. Mashour GA, Roelfsema P, Changeux JP, Dehaene S. Conscious Processing and the Global Neuronal Workspace Hypothesis. Neuron. 2020;105(5):776–98. doi: 10.1016/j.neuron.2020.01.026. PubMed PMID: 32135090; PubMed Central PMCID: PMCPMC8770991.

56. Minxha J, Mamelak AN, Rutishauser U. Surgical and Electrophysiological Techniques for Single-Neuron Recordings in Human Epilepsy Patients. Neuromethods. 2018;134:267–93. doi: 10.1007/978-1-4939-7549-5_14. PubMed PMID: WOS:000432982900016.

57. Behrens TE, Berg HJ, Jbabdi S, Rushworth MF, Woolrich MW. Probabilistic diffusion tractography with multiple fibre orientations: What can we gain? Neuroimage. 2007;34(1):144–55. Epub 20061027. doi: 10.1016/j.neuroimage.2006.09.018. PubMed PMID: 17070705; PubMed Central PMCID: PMCPMC7116582.

58. Rosen BQ, Halgren E. A Whole-Cortex Probabilistic Diffusion Tractography Connectome. eNeuro. 2021;8(1). doi: 10.1523/ENEURO.0416-20.2020. PubMed Central PMCID: PMCPMC7920542.

59. Glasser MF, Coalson TS, Robinson EC, Hacker CD, Harwell J, Yacoub E, et al. A multi-modal parcellation of human cerebral cortex. Nature. 2016;536(7615):171–8. Epub 20160720. doi: 10.1038/nature18933. PubMed PMID: 27437579; PubMed Central PMCID: PMCPMC4990127.

60. Delorme A, Makeig S. EEGLAB: an open source toolbox for analysis of single-trial EEG dynamics including independent component analysis. J Neurosci Meth. 2004;134(1):9–21. doi: 10.1016/j.jneumeth.2003.10.009. PubMed PMID: WOS:000189330500002.

61. Zanos TP, Mineault PJ, Pack CC. Removal of spurious correlations between spikes and local field potentials. J Neurophysiol. 2011;105(1):474–86. doi: 10.1152/jn.00642.2010.

62. Schomburg EW, Anastassiou CA, Buzsaki G, Koch C. The spiking component of oscillatory extracellular potentials in the rat hippocampus. J Neurosci. 2012;32(34):11798–811. doi: 10.1523/JNEUROSCI.0656-12.2012. PubMed PMID: 22915121; PubMed Central PMCID: PMCPMC3459239.

