## supplementary-figures for "Cross-region neuron co-firing mediated by ripple oscillations supports distributed working memory representations"

**Competing interests:** The authors declare no competing interests.

**Acknowledgements:**
We thank Sierra Wilson, Adam Niese, and Jacob Garrett for their support. This work was supported by NIMH (T32 MH020002, F31 MH135645). Acquisition, processing, and public release of the data was supported by the NIH BRAIN initiative through U01NS117839.

### Supplementary Figures

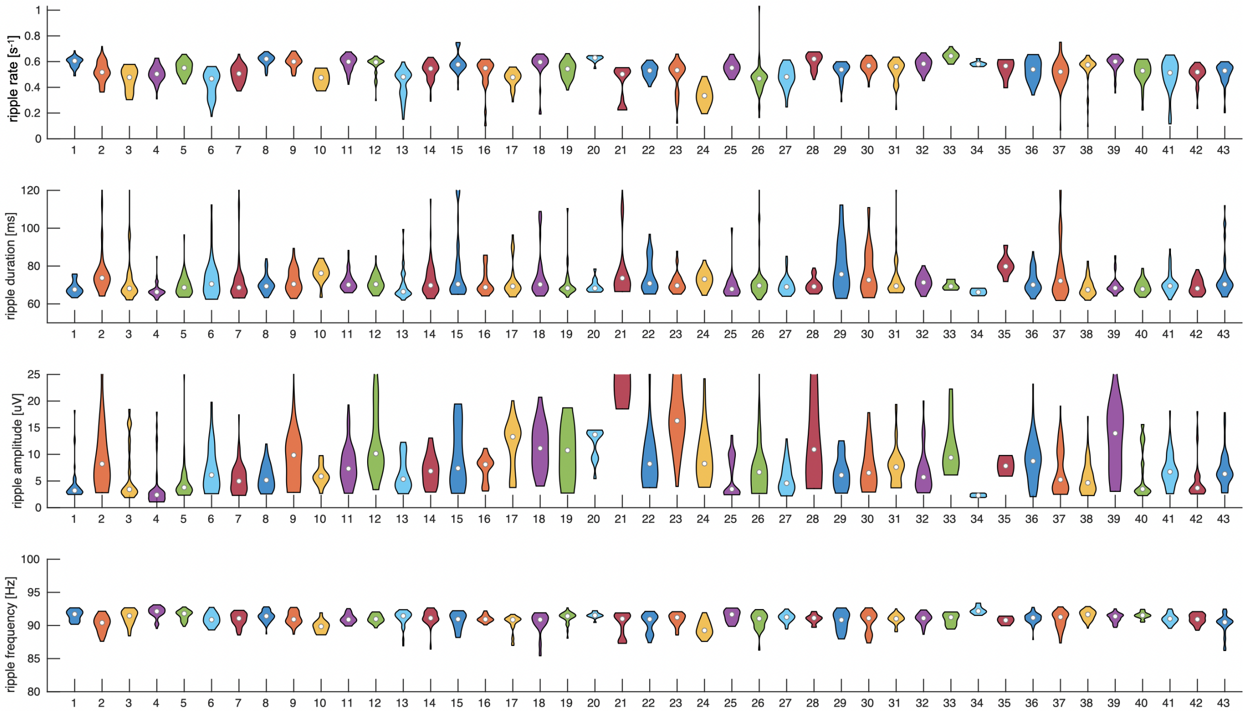


**Supplementary Figure 1 - Distribution of per-channel ripple characteristics separated by recording session**. Data is plotted for **ripple** event rate, duration, amplitude and oscillation frequency. Circle shows median and violin plots show the underlying distribution**.** Session number is shown on the x-axis (43 sessions across 35 patients)

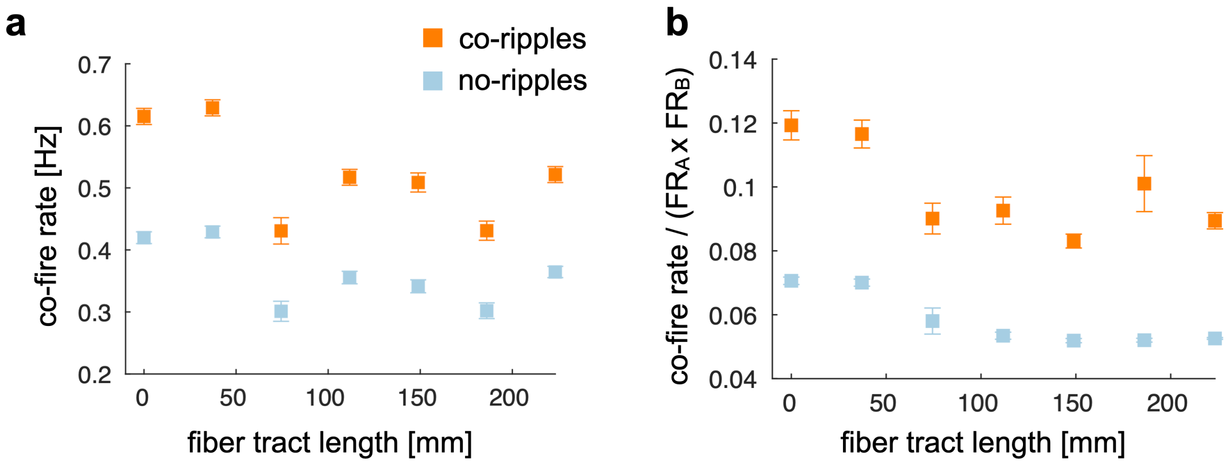


**Supplementary Figure 2 – Co-firing trends between units as a function of white matter fiber tract distance** Data is plotted across 7 equally spaced bins for co-firing rate **(a)** and co-firing normalized by the product of the baseline firing rate for each cells **(b)**. Co-firing is defined with a $\Delta t$ of ± 25 ms.

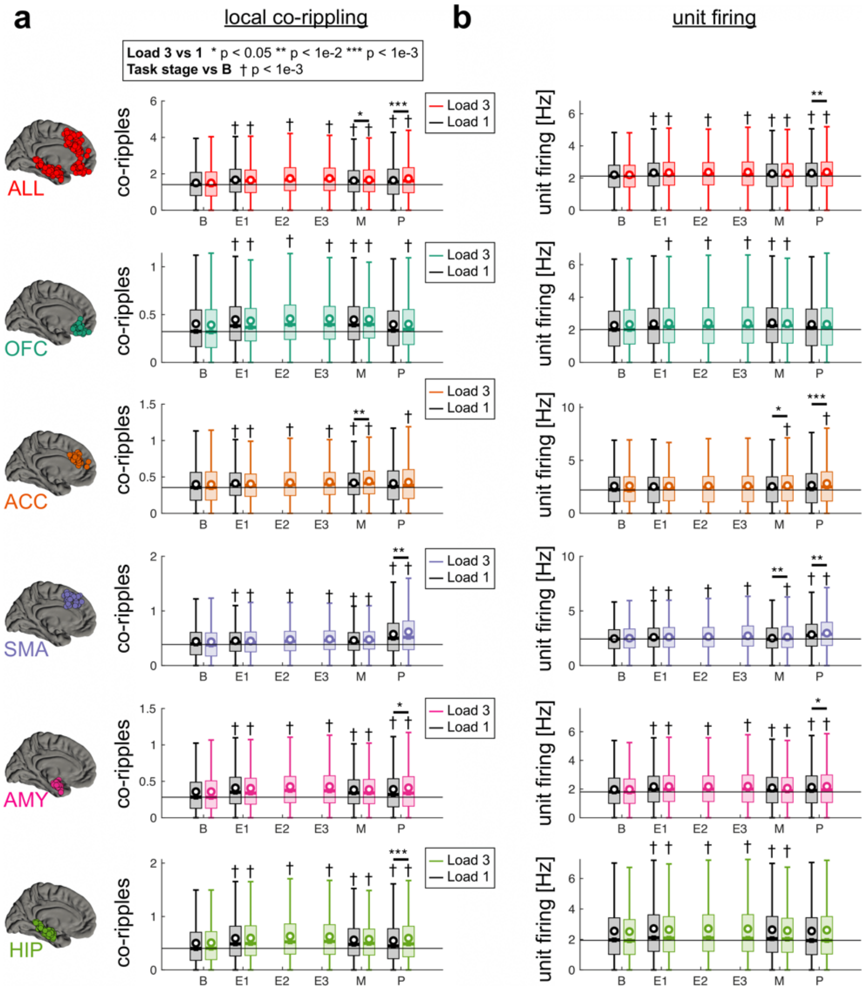


**Supplementary Figure 3 - Co-ripple and unit firing distribution for load 1 and load 3 in each stage of the task**. **(a)** Load 3 trials are shown in color and load 1 in black. * p<0.05, ** p<1e-3, *** p<1e-4 load 3 vs load 1 linear mixed effects FDR-corrected. † p<0.05 load vs baseline (B) FDR-corrected. **(b)** same as **(a)** but for single neuron firing rate within each region. Circles show mean, boxes show median and inter-quartile range and whiskers show 95%ile. Differences between load are tested with linear mixed effects modeling and differences from baseline are tested with two-sided Wilcoxon rank sum test.

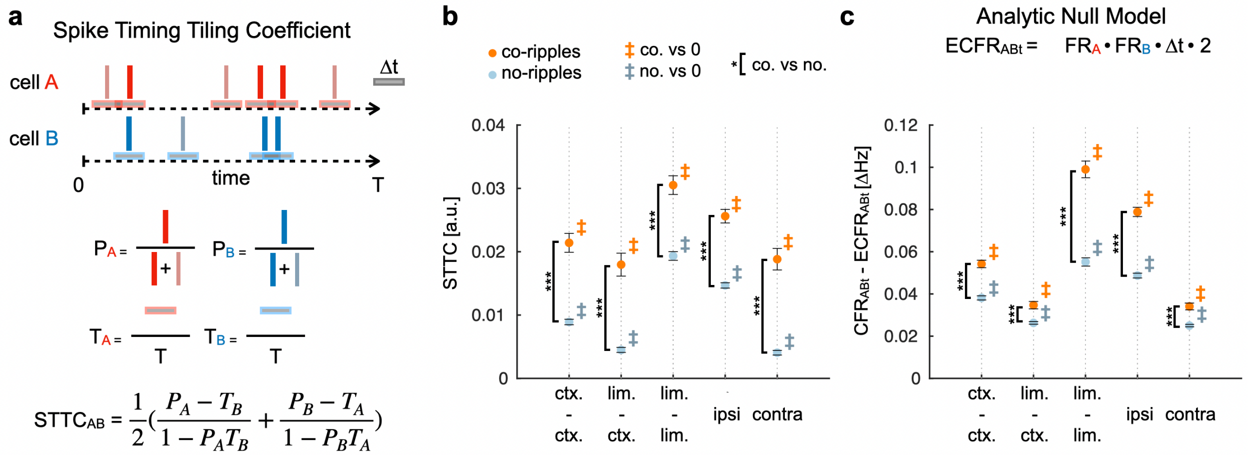


**Supplementary Figure 4**. Co-ripple enhancement of pairwise neural synchrony beyond expectations from cell firing rates, as calculated using two analytic methods. (a) Schematic illustrating the STTC calculation method. For two cells (A and B), spikes occurring within a time window (±Δt) of spikes from the other cell are identified. $P_A$ and $P_B$ represent the proportion of spikes from each cell falling within ±Δt of any spike from the other cell; $T_A$ and $T_B$ represent the proportion of total recording time covered by ±Δt windows around each cell's spikes. STTC is computed as the average of the two normalized terms, yielding values bounded between -1 and +1. (b) STTC values comparing co-ripple (orange) versus no-ripple (blue) periods across different regional pairings: cortical-cortical (ctx.-ctx.), limbic-cortical (lim.-ctx.), limbic-limbic (lim.-lim.), ipsilateral (ipsi), and contralateral (contra) pairs. Cell pairs are selected if they contain at least one co-firing event during co-ripples and no-ripples. Both co-ripple and no-ripple periods show STTC significantly greater than zero (‡), but STTC during co-ripples is significantly elevated compared to no-ripples across all connection types (*). (c) Observed co-firing rate of cell-pairs above the estimated co-firing rate calculated using a simple analytic model assuming that the spikes of A and B are renewal processes and completely unrelated. Estimated minus observed co-firing is calculated separately during co-ripple versus no-ripple periods. $FR_A$, $FR_B$ = measured firing rates of cells A and B; $CFR_{ABt}$ = measured co-firing rate of A and B within ±Δt; $ECFR_{ABt}$ = $FR_A$ x $FR_B$ x Δt x 2 = estimated co-firing rate of A and B within ±Δt, assuming that the spikes of A and B are renewal processes and completely unrelated. Co-firing exceeds the rate-corrected null during both conditions (‡), but co-ripples show significantly greater elevation than no-ripples across all connection types (*). Error bars represent SEM. ‡ $p < 0.001$ vs. 0; *$p < 0.001$ co-ripples vs. no-ripples.

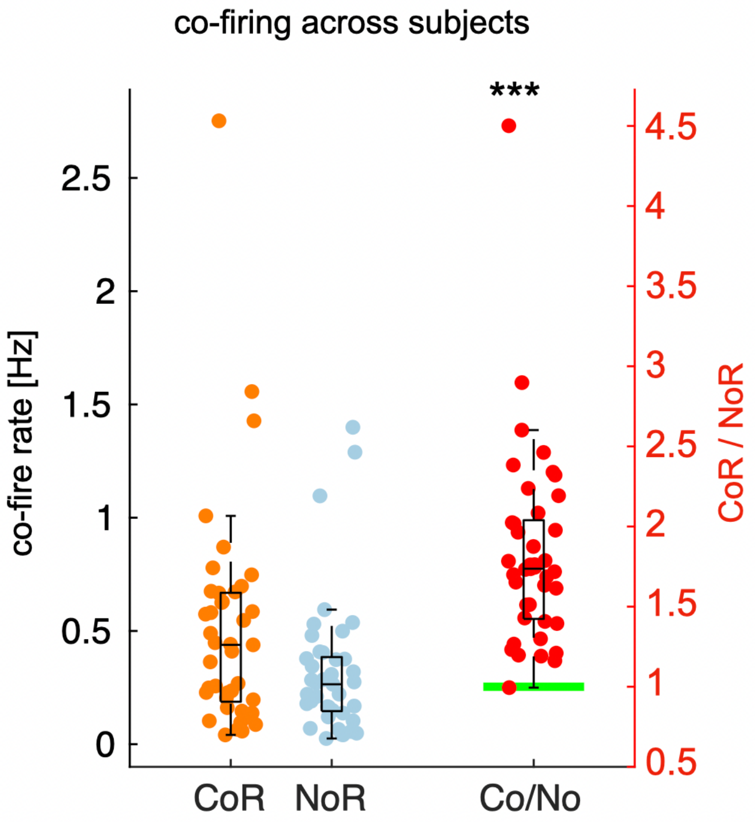


**Supplementary Figure 5. Co-firing rates are consistently elevated during co-ripple events across subjects.** Co-firing rates (left y-axis, Hz) during co-ripple (CoR, orange) and no-ripple (NoR, blue) periods, with individual data points representing individual subjects. The ratio of co-firing during co-ripples relative to no-ripples (Co/No, red; right y-axis) demonstrates that the majority of subjects exhibit enhanced co-firing during co-ripple events, with values predominantly exceeding 1 (green reference line). Box plots display median and interquartile range; whiskers extend to data within 1.5× IQR. ***p < 0.001, permutation test of medians.

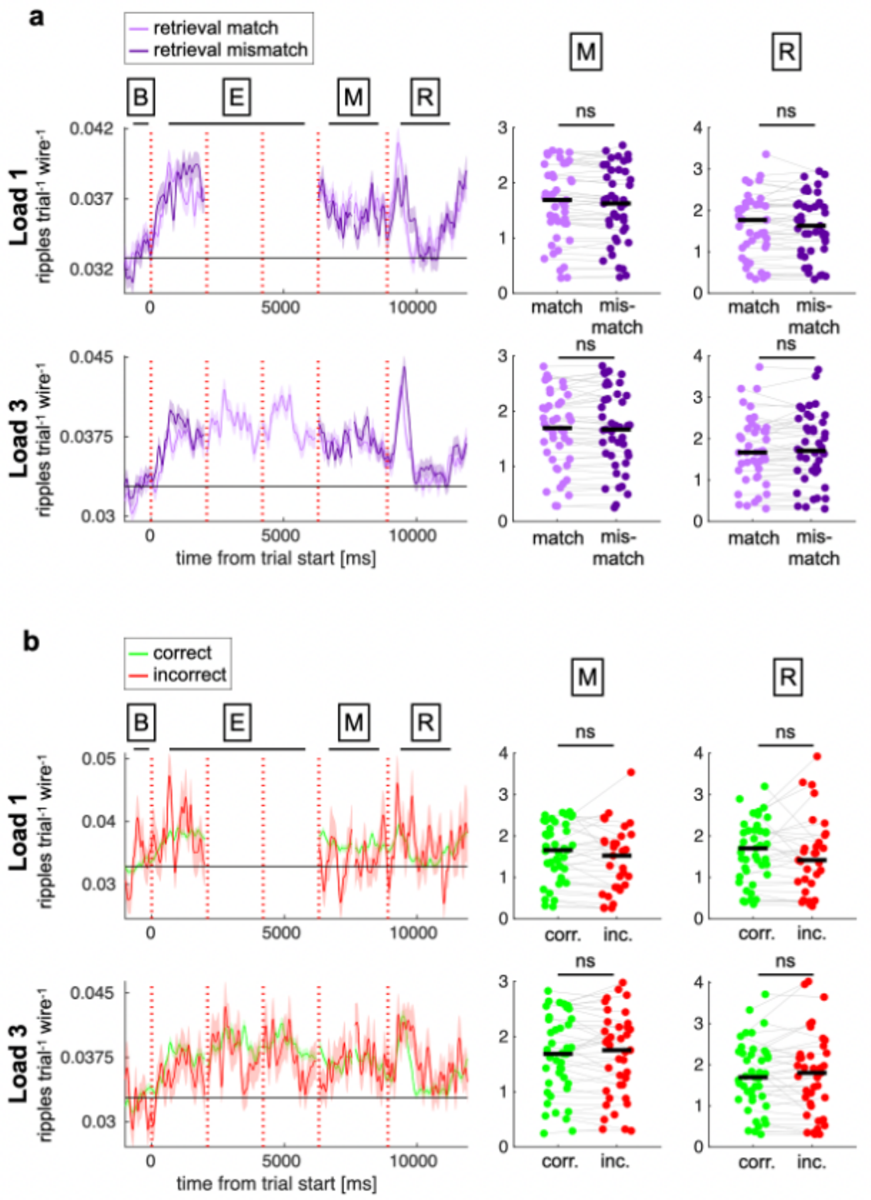


**Supplementary Figure 6 - Ripple rates do not robustly differentiate match/mismatch or correct/incorrect trials. (a)** Ripple rates comparing retrieval match (light purple) versus mismatch (dark purple) trials during load 1 (top) and load 3 (bottom) conditions. Left panels show ripple density (ripples per trial per wire) across task phases: baseline (B), encoding (E), maintenance (M), and retrieval (R). Vertical dashed lines indicate task phase boundaries. Right panels show subject-level ripple rates during maintenance and retrieval periods; gray lines connect paired observations. Black horizontal lines indicate means. **(b)** Ripple rates comparing correct (green) versus incorrect (red) trials during load 1 (top) and load 3 (bottom) conditions, plotted as in (a). While ripple rates increase during task periods relative to baseline across all conditions, neither match/mismatch status nor behavioral accuracy robustly modulates ripple density. *p < 0.05, ns = not significant, permutation test of means.

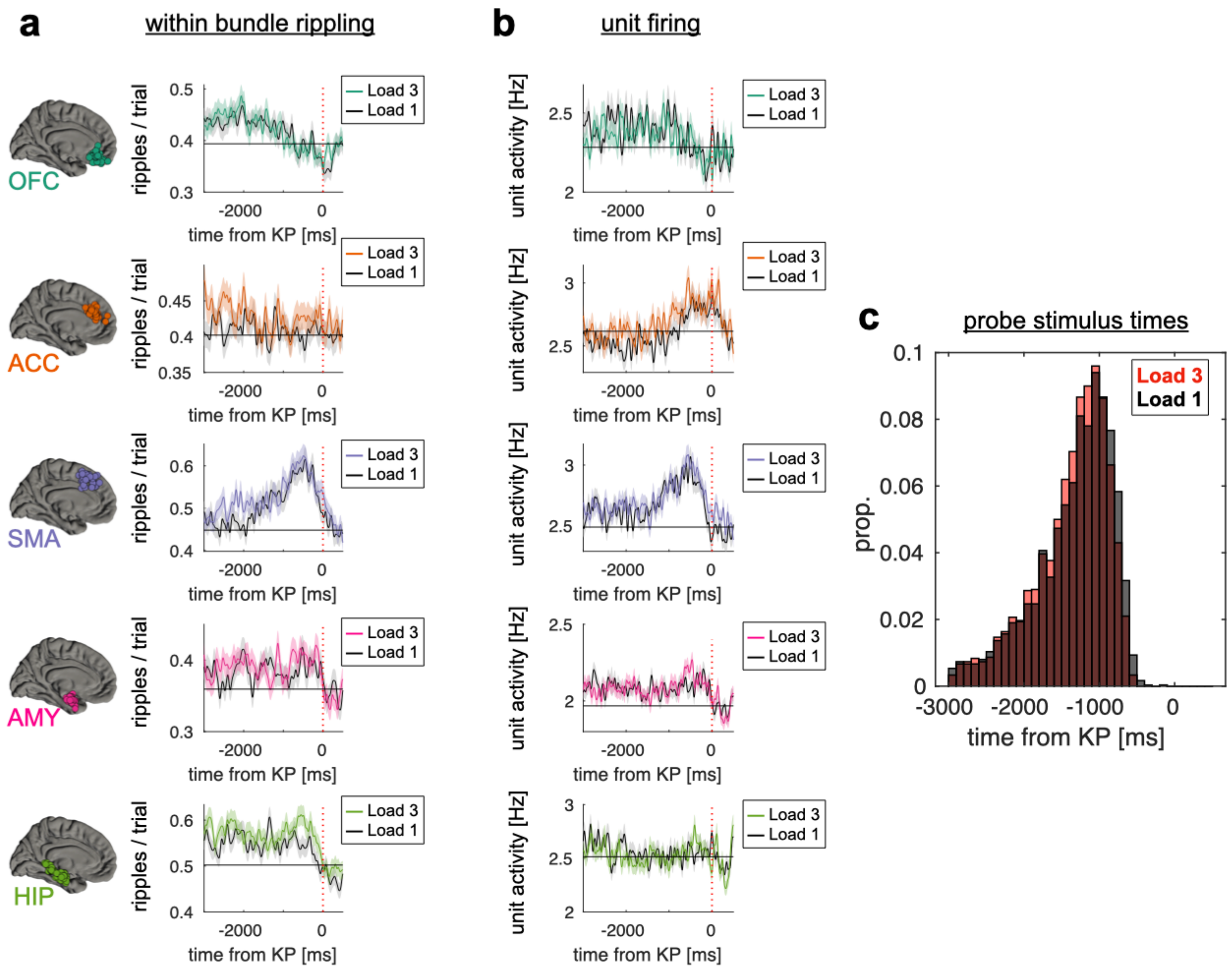


**Supplementary Figure 7 – Ripple rates locked to response keypress**. Same as Fig. 2, but locked to response keypress (KP) for **(a)** ripple oscillation rates and **(b)** unit firing within each sampled region. Data is shown for mean number of rippling electrodes across all trials. Colored curves show mean ± sem for all load 3 trials, while black curves show load 1 trials. For plotting purposes, data are smoothed with a 100 ms sliding gaussian window. (c) Histogram of probe stimulus onset times relative to KP for load 1 and 3.

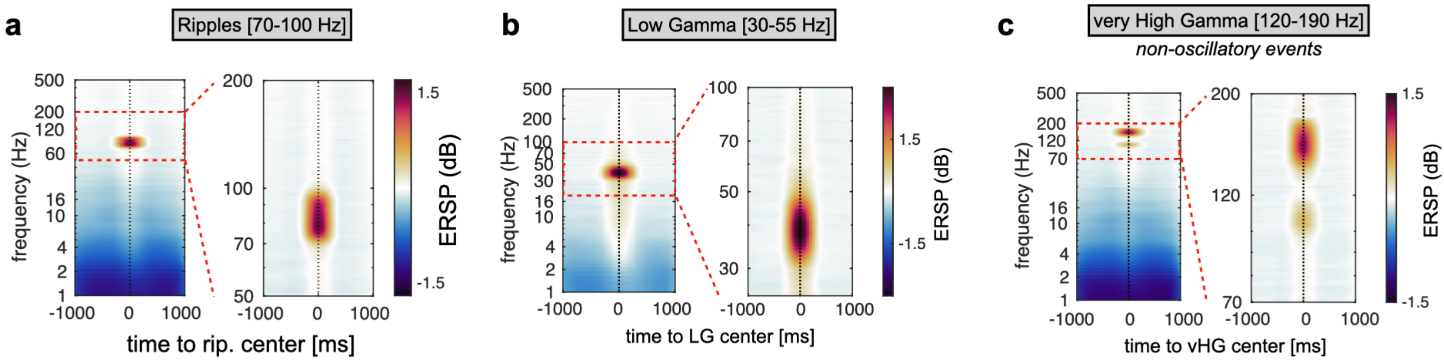


**Supplementary Figure 8** . Average time-frequency plot across all microelectrodes locked to event centers (left), and zoomed in around the event frequency range (right). Data is shown for ripples **(a)**, low gamma oscillations **(b)**, and non-oscillatory very high gamma **(c)**.

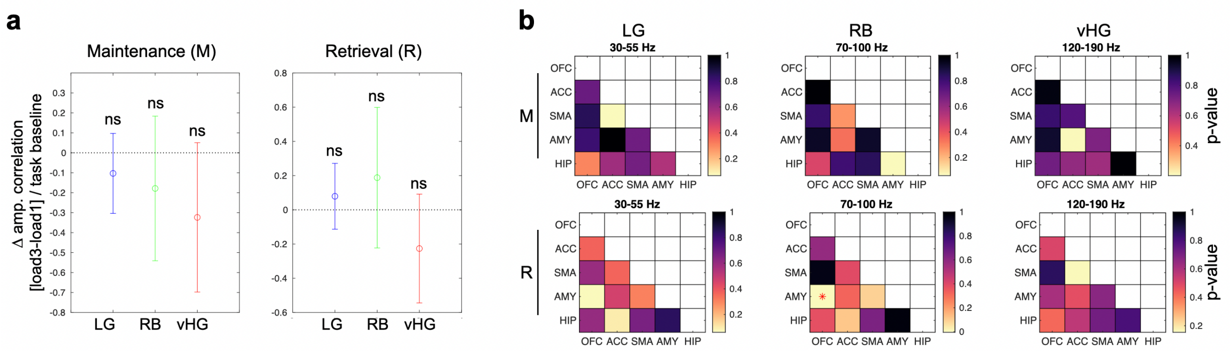


**Supplementary Figure 9. Cross-regional amplitude envelope correlations do not robustly scale with memory load. (a)** Change in amplitude envelope correlation between load 3 and load 1 conditions, normalized to task baseline, during maintenance (M, left) and retrieval (R, right) periods across three frequency bands: low gamma (LG, 30–55 Hz), ripple band (RB, 70–100 Hz), and very high gamma (vHG, 120–190 Hz). Error bars represent SEM. ns = not significant. **(b)** Region-by-region p-value matrices for load-dependent changes in amplitude correlation during maintenance (M, top row) and retrieval (R, bottom row) across frequency bands. Regions include orbitofrontal cortex (OFC), anterior cingulate cortex (ACC), supplementary motor area (SMA), amygdala (AMY), and hippocampus (HIP). Warmer colors indicate lower p-values. Asterisk denotes $p < 0.05$ .

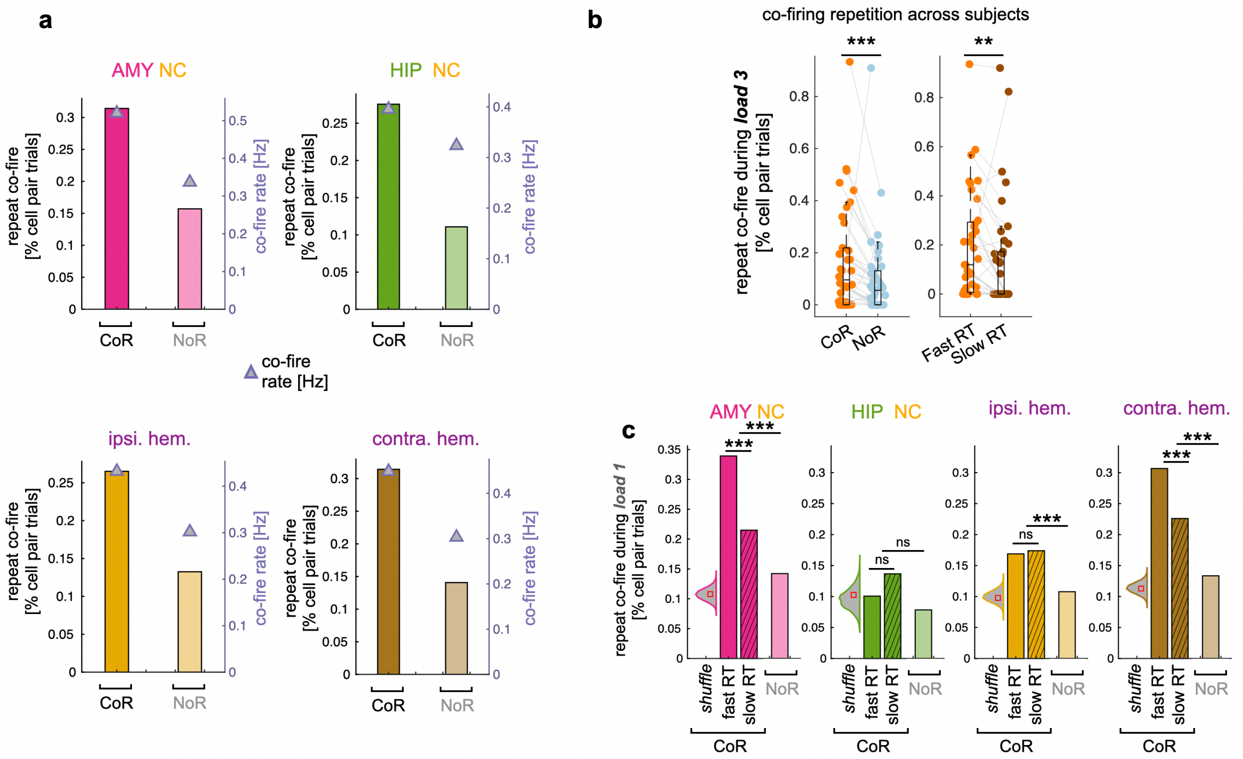


**Supplementary Figure 10. Co-firing repetition during co-ripples across subjects, regional pairings, and additional memory load conditions. (a)** Repeated co-firing between encoding and retrieval (% cell pair trials, bars; left y-axis) and co-firing rate (Hz, triangles; right y-axis) during co-ripple (CoR) versus matched no-ripple (NoR) periods across regional pairings: amygdala-neocortex (AMY NC), hippocampus-neocortex (HIP NC), ipsilateral hemisphere (ipsi. hem.), and contralateral hemisphere (contra. hem.). Co-firing repetition is elevated during co-ripples compared to no-ripple periods across all connection types, even when accounting for overall increases in co-firing rate. **(b)** *Subject-level distribution of repeated co-firing during load 3 (highest memory demand) trials, expressed as percentage of cell pair trials.* ***(Left)*** *Co-ripple (CoR, orange) versus no-ripple (NoR, blue) periods.* ***(Right)*** *Fast response time (Fast RT, orange) versus slow response time (Slow RT, brown) trials. Each data point represents an individual subject; gray lines connect paired observations. The majority of subjects show elevated co-firing repetition during co-ripple events and during trials with faster behavioral responses. Box plots display median and interquartile range; whiskers extend to data within 1.5× IQR. **p < 0.01, ***p < 0.001, paired permutation test of medians.* **(c)** Repeated co-firing during load 1 (lowest memory demand) trials across regional pairings. Bars show co-ripple periods stratified by behavioral performance (shuffle control, fast RT, slow RT) and no-ripple periods. Violin plots display the distribution of shuffle controls. Open squares indicate shuffle medians. Co-firing repetition effects during co-ripples remain significant even under low memory load, though the relationship with reaction time is attenuated compared to high load conditions. Statistics performed using chi squared. **p < 0.01, ***p < 0.001, ns = not significant.
